# Visual Word Form Area demonstrates individual and task-agnostic consistency but inter-individual variability

**DOI:** 10.1101/2025.07.23.666206

**Authors:** Jamie L. Mitchell, Mia Jimenez, Hannah L. Stone, Maya Yablonski, Jason D. Yeatman

## Abstract

Ventral Occipital Temporal Cortex (VOTC) contains a mosaic of categorically-selective functional regions that respond to visual stimuli. Within left VOTC lies the Visual Word Form Area (VWFA) - a text-selective region that develops as individuals learn to read. While there is consistency in the general location of text-selective responses - within the posterior occipitotemporal sulcus (OTS) - there is substantial variability across individuals in the size and precise anatomical location. Moreover, debate continues regarding whether VWFA a) encodes the visual features of text, versus b) is driven by the task of reading. Using functional magnetic resonance imaging, we scanned adults and children as they completed two tasks while viewing text, pseudo fonts, faces, objects, limbs: (1) a fixation task where participants ignored stimuli while making psychophysical judgements on a fixation dot, and (2) a one-back task, where they attend to stimuli to detect repeats. We found a consistent VWFA location could be identified on each individual’s cortical surface using either task. At the same time, the one-back task evoked a larger territory of text-selective response (leading to a larger ROI) than the fixation task. However, when averaged in template space, text-selective cortex could not be identified due to variability in the relative locations of text-, face-, object-, and limb-selective cortex. Thus, despite task-agnostic consistency in individual VWFA, text-selective responses are masked when averaged in template space due to variability in the exact configuration of category-selective regions. Furthermore, these effects were present in both children and adults.

## 1 Introduction

Deriving meaning from symbols on a page relies on the successful processing of visual stimuli before visual information can be conveyed to language regions of the brain^1^. One brain area commonly studied in reading development is a region of visual cortex, known as the Visual Word Form Area (VWFA), that is tuned to the visual features that comprise a person’s native orthography^2,3^. This text-selective patch of cortex, which develops in the posterior and lateral portion of the left ventral occipitotemporal cortex (VOTC), is generally localized to the occipital temporal sulcus (OTS)^4,5^. VWFA is a pivotal neural structure for proficient reading and is hypothesized to be instrumental in the early word recognition, identifying words and letters from lower-level visual shapes before their association with phonology or semantics^1,6^. However, since the original report of a “visual word form area” localized to high-level visual cortex based on fMRI^7^, there has been a debate over whether this region of cortex a) encodes the visual features of text versus b) is driven by the task of reading (or language more broadly) and should not be considered a visual region per se^8–11^. A primary reason this debate has persisted is due to methodological inconsistencies in the experiments that have been used to study the VWFA as well as the analytic approaches that are used to define this region.

Inconsistencies exist regarding the proper way to isolate VWFA. Previous studies suggest that the VWFA typically emerges with peak activation around MNI coordinates x = -43, y = -54, z = -12^6^, suggesting a robust anatomical anchor for the region. While functional localizers can successfully isolate a patch of cortex that selectivity responds to text on the individual cortical surface^4,5,12^, it is important to distinguish this from the claim of its functional consistency across different tasks. Although a region can be reliably found with a specific localizer, its precise boundaries might shift, and its activity profile can be dynamically modulated depending on the specific task demands or stimuli presented. Some have claimed that task, rather than stimulus, is the primary factor driving localization implying that there is no consistent text-selective patch of cortex^11,13–15^.

Despite broad consensus on the VWFA’s involvement in reading, fundamental questions persist concerning its exact functional role^10,16^, its degree of specificity^11^, and the intricate neural circuitry that supports its computations^5^. A central point of contention revolves around whether VWFA exclusively processes written words or if its activity extends to other visual categories and is primarily driven by task demands^10,17,18^. Fiez and Petersen^19^ noted that the use of various tasks to study the same region leads to variability among results. Others argue that the “purported VWFA” is actually the result of top-down signals from language regions and has nothing to do with encoding the visual features of text^11,13–15^. This challenge is compounded by findings in lower-level visual cortex, where tasks and arousal states significantly influence blood-oxygen-level-dependent (BOLD) response^20,21^. While the VWFA displays inherent orthographic selectivity, its functional profile adapts under top-down control from the language network as task demands shift.^22,23^. This dynamic modulation of functional characteristics, rather than a fixed, static response, contributes to the complexity in fully characterizing the anatomy and function of the VWFA.

A significant contributor to conflicting findings regarding VWFA specificity is the common practice of employing group-level analyses in template space (i.e., MNI^24^ or fsaverage^25^) or template-based ROIs^26^. Aligning individual participants to an “average brain” smears BOLD signals across cortical areas, including non-word-selective regions, since the VWFA is located slightly differently in each individual. This directly explains why some studies, particularly those relying on group-level analyses in template space, might report less specificity (or even non-existence) for the VWFA^16,27,28^, while others utilizing individual localizers demonstrate strong word-selectivity^26,29^. Consequently, template ROIs might inadequately analyze individual response profiles as they fail to precisely identify the text-selective cortex within individuals.

Thus signals are contaminated by neighboring regions with dramatically different selectivity patterns. In other words, an area falling within a template ROI for one person may fall outside it for another. The inherent variability in the precise anatomical location and size of the VWFA across individuals means that averaging functional data across participants can effectively “wash out” the selectivity of the region and give the appearance that it responds to other stimuli (e.g., faces or objects) or is driven by various tasks.

VOTC houses regions that selectively respond to specific categories of visual images including faces, places, bodies and, of course, text^4^. Some argue that object recognition is the result of distributed activation patterns throughout this broad swath of cortex^18^, while others argue that discrete patches of cortex within VOTC are responsible for recognition of specific categories^30^. While there is still debate about the modularity of VOTC, it is well documented that VWFAs emerge developmentally later than other categorically-selective regions^31,32^. As individuals learn to read, dynamic changes unfold within VOTC resulting in the emergence of a VWFA. Reading, an evolutionarily recent skill, has not afforded the brain adequate time to develop innate regions for visual word recognition. In contrast to evolutionarily ancient categories, like faces processed in the Fusiform Face Area (FFA), text processing must necessarily leverage and repurpose pre-existing neural circuitry. The neuronal recycling hypothesis^33^ suggests that reading acquisition recycles or repurposes a region of the ventral visual cortex initially involved in more general object recognition or other visual processing^34–36^. Further evidence suggests that the development of the Visual Word Form Area is closely tied to reading ability and literacy acquisition^29,32^. The very mechanism of this recycling provides a fundamental explanation for individual variability in the VWFA. The precise pre-existing neural biases and the specific “path” of this recycling process may differ slightly across individuals, leading to observable variations in the VWFA’s size, exact anatomical location, and functional tuning. This contrasts with more evolutionarily ancient systems, which might exhibit less inter-individual variability in their core functional organization, thereby supporting the assertion that text processing represents a unique case in functional organization.

While VWFA is consistently localized across individuals and writing systems^37^, a significant degree of variability in its size and precise anatomical location is also well-documented; likely due to its unique experience-dependent developmental trajectory. As noted by Glezer and Riesenhuber^26^, obscuring of individual characteristics through group-level approaches leads to the very controversies surrounding VWFA that is observed in the literature, yet this practice is still widely utilized over a decade after their initial observation. Despite the extensive research on the VWFA, critical gaps remain in our understanding of its variability across individuals and its functional characteristics under different task demands. Prior studies, including those by Glezer and Riesenhuber, have employed group-level analyses, which obscure the nuances of individual differences in VWFA localization and functioning.

Furthermore, the implications of these individual differences for our understanding of the developmental trajectory of reading skills and the potential influence of varying task demands have not been sufficiently explored. By expanding on Glezer and Riesenhuber’s work, this study aims to illuminate the complexities of the VWFA and its neighboring visual regions, ultimately contributing to a more nuanced understanding of the neural basis of reading. In this study, we expand on Glezer and Riesenhuber’s work and test two specific hypotheses regarding the consistency and variability of the VWFA and surrounding regions:

Hypothesis 1: Within an individual, there is a consistent location within VOTC that responds to text, irrespective of the task demands.

Hypothesis 2: The precise configuration of this text-selective region relative to face-, object-, and limb-selective regions will vary across individuals, such that group averages in template space will mix signals among multiple regions leading to misleading results.

We address these hypotheses in two separate samples: (1) a diverse and representative sample of children with variable reading ability and (2) a homogenous sample of literate adults who are skilled readers. Participants were scanned while engaging in different tasks involving different visual categories, including real words, pseudo words, faces, objects, and limbs. By adopting an individual-level analysis approach, categorically-selective ROIs were defined on each individual’s native cortical surface. We expand on the approach of Glezer and Riesenhuber, examining metrics of size, task effect, and text-selectivity to further investigate the ways methodological choices can have major effects of the study of the reading circuitry and explore how these effects are present in both in a pediatric and adult population.

## 2 Methods

### 2.1 Participants

To understand how the anatomical and varying functional landscape affects group-level analyses of the VWFA, 82 children (ages 7-13; 39 female) and 14 adults (ages 21-34; 6 female) underwent functional magnetic resonance imaging (fMRI) scans. Child data was collected as part of a larger longitudinal study^29,38^. The children in this study came from a highly diverse sample spanning a large range of socioeconomic status, reading ability, language, ethnic and racial backgrounds, and educational experiences. As such, the child sample in this study is a very heterogeneous sample representative of the population in the broad San Francisco Bay Area. Adult participants were recruited from a convenience sample around Stanford University campus and consisted primarily of university students, representing a more homogenous sample. A total of 87 children and 15 adults were recruited, however data for 5 children and 1 adult were not included in the current analysis (see Data Inclusion for more details).

All adult participants gave written informed consent and child participants gave informed verbal assent accompanied with parent informed written consent in compliance with the Stanford University Institutional Review Board. Participants all reported normal or corrected-to-normal vision and self-identified as either a monolingual English speaker or as using English for at least 60% of their daily communication. See Table 1 for more detail on participant demographic information.

**Table 1.**
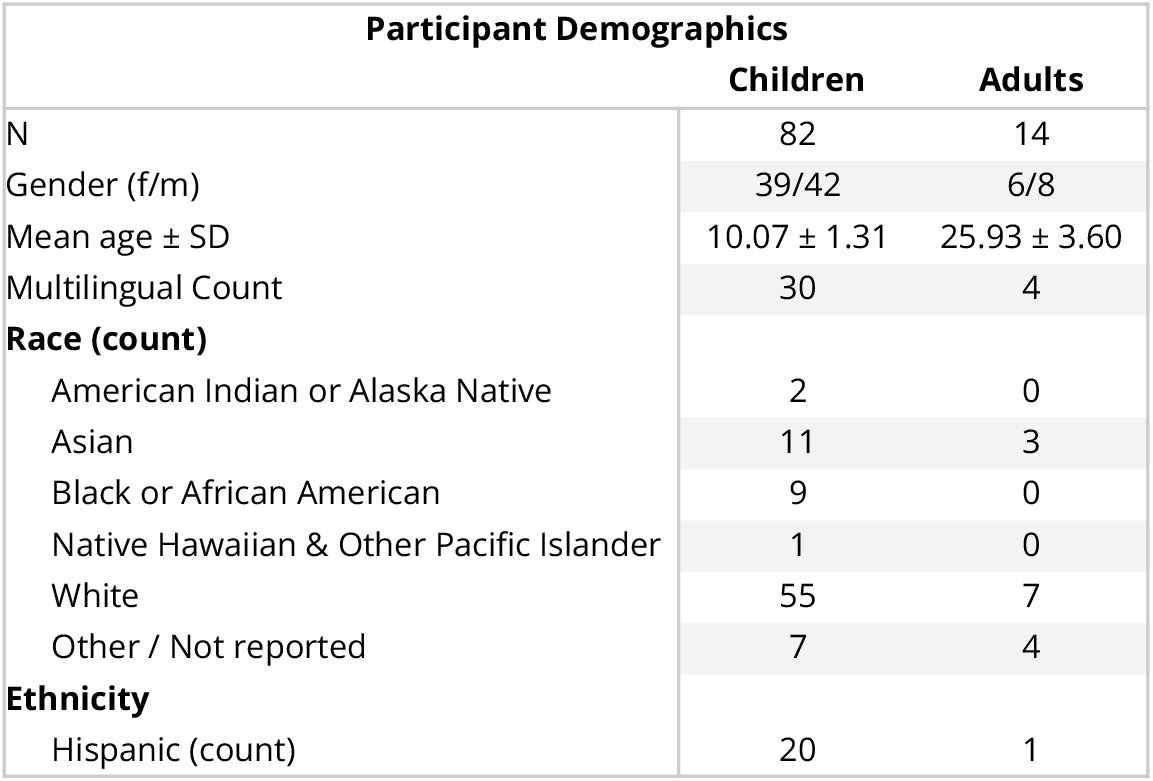
Participant Information. Demographic information separated by age group. Gender, multilingual status (speaking more than one language on a daily basis), race, and ethnicity are collected as self-report responses.

| Participant Demographics |  |  |
| --- | --- | --- |
|  | Children | Adults |
| N | 82 | 14 |
| Gender (f/m) | 39/42 | 6/8 |
| Mean age ± SD | 10.07 ± 1.31 | 25.93 ± 3.60 |
| Multilingual Count | 30 | 4 |
| **Race (count)** |  |  |
| American Indian or Alaska Native | 2 | 0 |
| Asian | 11 | 3 |
| Black or African American | 9 | 0 |
| Native Hawaiian & Other Pacific Islander | 1 | 0 |
| White | 55 | 7 |
| Other / Not reported | 7 | 4 |
| **Ethnicity** |  |  |
| Hispanic (count) | 20 | 1 |

### 2.2 Experimental Design

The experiment consisted of two runs of two separate functional localizer tasks (one-back and fixation) in which participants were shown several images of different types of visual stimuli; an experiment that largely followed the localizer design described White and collegues^23^. In both tasks, participants were asked to fixate on a colored dot at the center of the screen. During the one-back task, participants were instructed to respond by pressing a button on a response box every time the image on the screen repeated. During the fixation task, participants were instructed to respond every time the fixation dot at the center of the screen changed color. In each run, images repeated and the fixation dot changed color at random in 33% of trials. Thus, the stimulus presentation and trial structure was kept identical for both tasks with the only difference being the instructions given to direct the task.

Each run of the experiment consisted of 65 4-second trials. Each trial contained a sequence of 4 images from the same stimulus category presented for 800 ms with 200 ms of blank fixation in between. Main categories of stimuli included text, pseudo fonts, objects, faces, and limbs. A “blank” category was also included and trial order was randomized. See Mitchell et al.^29^ and Stone et al.^38^ for more information on the experiment.

### 2.3 MRI Acquisition

Structural and functional images were collected at the Stanford Center for Cognitive and Neurobiological Imaging (CNI) on a 3 T GE Discovery MR750 fitted with a Nova-Medical 32-channel head coil. A whole-brain anatomical scan was acquired using a T1-weighted magnetization-prepared rapid gradient echo sequence with a 0.9 mm^3^ resolution. Functional data was collected using four runs of an echo-planar imaging sequence (2.4 mm^3^ voxels, repetition time of 1.19 seconds, echo time of 30 ms, flip angle of 62°, field of view of 20.2°, with 51 slices collected with a multiband of 3). Participants viewed the screen via a series of mirrors mounted to the head coil on the scanner reflecting onto a 47” 3D LCD Resonance Technology display mounted at the back of the magnet bore with a visual distance of 256.1 cm from the first mirror and a total of 277.1 cm from the participant’s eye.

To maximize data quality and minimize potential confounds, we monitored 1) motion in real time via FIRMM Real-Time Motion Tracking, 2) participant engagement and focus through a webcam3) responses during the task, and 4) potential artifacts in the fMRI data. If any data quality issues were detected (e.g., the participant closing their eyes, moving, etc.), the scan was ended early, and researchers communicated with the participant to ensure attentiveness before restarting the scan.

### 2.4 MRI Data Preprocessing and BOLD Response Estimation

Data preprocessing and blood oxygen level dependent (BOLD) signal calculation were completed using the pipeline outlined in Mitchell et al.^29^ and Stone et al.^38^. In short, we used fMRIprep^39^ to preprocess functional data, incorporating anatomical data aligned to the AC-PC axis and processed via FreeSurfer for segmentation and surface reconstruction. BOLD runs underwent various corrections, including head motion estimation, slice-time correction, and co-registration to the anatomical reference. The resulting BOLD time-series were resampled into surface space average responses were estimated using a general linear model that incorporated confound regressors, providing results in percent signal change relative to blank trials. T1w images used for preprocessing and visualizations were taken from the first time point where a high-quality image was successfully acquired from each child. Adult participants underwent identical preprocessing procedures.

### 2.5 Data Inclusion

87 children were originally recruited and participated in a larger longitudinal project. For the present study, data from a single time-point was used for analysis. Participants completed 4 runs of a functional localizer at every visit and the first time-point in which a child participant successfully completed all four runs of the experiment with high data quality (described below) was used for the current study. Adults completed a single visit and again participants who completed all four runs of the experiment with high data quality were included in the present study. Participants were excluded from the study if they 1) fell asleep or failed to keep their eyes open during any run of the experiment 2) had a mean framewise displacement (FD) of 0.5 mm or greater on any given run 3) if more than 30% of volumes had a FD of 0.5 mm or greater 4) did not complete all four runs of the experiment (two runs of the fixation task and two runs of the one-back task). After accounting for inclusion criteria, we were left with 82 children and 14 adults in our final study sample. The final sample of child participants had a mean FD of 0.186 (sd = 0.0889) and mean tSNR of 56.644 (sd = 11.901), and the final adult sample had a mean FD of 0.095 (sd = 0.035) and mean tSNR of 62.101 (sd = 6.682).

### 2.6 Categorical Response Estimation

To define functional regions of interest (ROIs) on each participant’s native surface, we fitted a general linear model (GLM) to the data using the Nilearn python package^40,41^. Beta estimates from the GLM were used to compute contrast maps to create surface meshes of category-specific responses for each participant. First, a GLM was fit to all four runs of the experiment to produce a single set of category-level responses separately within each participant. Using these estimates, three types of activation maps were created: 1) t-statistic maps for ROI definition, created by comparing activation to each category relative to the weighted average of all other categories ( i.e. Text > Other), 2) contrast maps for group-level modeling, created by taking the contrast values from the above contrast maps, 3). Activation or percent signal change maps for calculating category-specific response, created by taking the contrast values from a comparison of each category relative to baseline (i.e. Text > Blank).

Next, to investigate category-selectivity for each task separately, the data were split in half and a second GLM was fit to the one-back task runs and the fixation-task runs separately to derive activation maps by task. Once again, t-statistic maps were derived in the same manner as before to define task-specific ROIs and activation maps were created and used to analyze task-specific activations.

### 2.7 Functional Regions of Interest

Functional ROIs were manually drawn for each participant using t-statistic maps on the Freesurfer^42^ Native surface. All ROIs were defined within the confines of each participants native ventral occipitotemporal cortex (VOTC), specifically located posterior to the anterior tip of the occipitotemporal sulcus (OTS), lateral to the collateral sulcus, anterior to the posterior transverse collateral sulcus, and medial to, as well as including, the OTS. T-statistic maps were thresholded at t > 3 and all ROIs were defined by grabbing any vertex that fell within specific anatomical boundaries which had a t-statistic at or above the threshold. VWFA ROIs were identified as all vertices that met the threshold value in a text > other contrast, and were located on the left occipitotemporal sulcus and the lateral region of the fusiform gyrus (any cortical area lateral to the mid-fusiform sulcus). FFA ROIs were defined as the vertices that met the threshold value in a faces > other contrast, and were found within the fusiform gyrus and mid-fusiform sulcus. Two sets of ROIs were defined with this criteria: First, VWFAs and FFAs were drawn using all four runs of data for each participant. Second, task-specific ROIs (denoted tVWFA and tFFA) were defined using the same approach for each participant, but using functional maps created for each task separately.

Group-level ROIs were also created to assess the degree of spatial overlap across participants. To do so, native ROIs were projected to fsaverage space and a probabilistic map was generated by calculating the overlap of each ROI and dividing by the total number of participants. These maps were thresholded at a probability of 0.2 (indicating more than 20% of participants had an ROI at a given vertex) to yield group ROIs^43,44^, labeled aVWFA and aFFA for adults, and cVWFA and cFFA for children.

Finally, literature-based functionally-defined ROIs were used for further analyses to compare effects of native and group labels. VWFAs and FFAs from Rosenke et al.^44^ and Kubota et al.^43^, defined with a similar probabilistic map approach, were used. For clarity, pOTS-characters ROI and the combination of the pFus-faces and mFus-faces ROIs from Rosenke and colleagues will be referred to in this paper as rVWFA and rFFA respectively. These ROIs were selected as they were defined from a probabilistic map in a sample of adult participants. Similarly, the combination of the pOTS-words and mOTS-words ROIs and the pFus-faces and mFus-faces ROIs from Kubota and colleagues will be referred to as kVWFA and kFFA respectively. These ROIs were again generated using a probabilistic approach, however the sample used to generated this template ROI included child participants. We ensured that template ROIs were selected as they were defined on age groups similar to those included in the present study.

### 2.8 Statistical Analysis

To investigate the text-selective responses in the ventral occipitotemporal cortex (VOTC), specifically focusing on the visual word form area (VWFA) and the fusiform face area (FFA), we employed a series of analyses to ensure robust identification and comparison of these regions across individuals. Unless specified, all analyses reported here consisted of two-sides statistical tests.

#### Consistency of VWFA Response Across Tasks

To evaluate the stability of the VWFA across different task demands, we divided our dataset by task (one-back vs. fixation). Each participant’s VWFA was defined independently using data from one task. We computed the Dice similarity coefficient (DSC) for the regions defined by each task to assess the overlap and consistency between these task-defined VWFAs. This was done with the following formula:

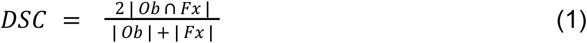

where *Ob* ∩ *Fx* is the total the number of vertices that intersect between both tVWFAs, and |*Ob* | + | *Fx*| is the sum of the total number of vertices in the surface mesh (which is twice the size of the participants native hemisphere).

#### Analysis of BOLD Responses

To analyze the BOLD responses within the defined ROIs, fit a linear mixed effects model (LME) using the lme4 package in R. We first estimated the response to each category (BOLD response) as a function of the two-way interactions between otVWFA/ftVWFA (ROI Task) & stimulus category (Category), and task-specific response (BOLD Task) & stimulus category with a random intercept of participant, treating the otVWFA and the one-back BOLD response to text as references in both children and adults, modeled with the following equation:

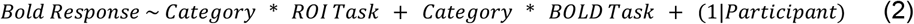

To determine how if and how the tasks effect individual category tuning different, we next fit another LME estimated BOLD response as a function of the interaction between the ROI task and the BOLD task. We again included a random intercept of participant and treated the one-back VWFA and the signal from the one-back task as the references, modeled with the following equation:

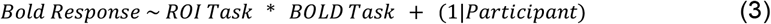

These models were designed to quantify the response magnitudes associated the visual stimuli and to identify any significant interaction effects relating to task demand.

#### Exploration of Text Selectivity

In addition to examining overall BOLD responses, we calculated a text selectivity index for each participant by comparing the response to text against other categories^29,32^. A separate LME, similar to equation 2, was fitted to assess this relationship:

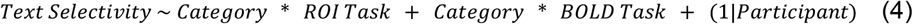

Through these models, we aimed to quantify how responsive VWFAs were to text compared to other visual categories across both one-back and fixation tasks.

#### Group-Level Analyses

To determine how the ROI used to analyze the tuning properties of the cortex, we conducted a series of analyses comparing response properties captured by the native and manually defines ROIs relative to each group and template ROI. To quantify the differences in response magnitude to each stimulus category, we fit a LME looking at BOLD responses as a function of ROI with a random intercept of participant and treated the mean response for native space ROIs as the reference for each stimulus category, again including a random intercept of participant. This was modeled as follows:

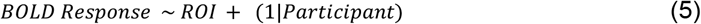

We then evaluated the selectivity of the various VWFAs with a similar approach as follows:

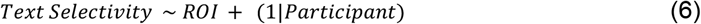

## 3 Results

### 3.1 Text-Selective Cortex Can Be Identified in Individual VOTCs

To assess how group level analyses may obscure individual level results related to VWFA, or other small category-selective regions dispersed among larger and more stable regions, we first ensured that VWFA exists on an individual level in our participants.To do so, we identified category-selective regions by contrasting neural response to each category against all other categories (for example, text versus the weighted sum of faces, objects, limbs, and pseudo-fonts, thresholded at t > 3) for every vertex in each hemisphere. ROIs were manually drawn on the cortical surface of each participant. Figure 1 shows this process with the projection of the Text > Other t-statistic map and both the VWFAs and FFAs of 6 sample participants (See Supplemental Figure S2 for the visualization of the Faces > Other map in the same individuals). Using this process, we were able to identify a VWFA in 69 child and 13 adult participants. This is unsurprising as this finding confirms what is already known from previous work which has shown that children only develop a VWFA as they attain a certain reading level^31,34,45^ and our child participants had a large range of reading ability^29^.

**Figure 1.**
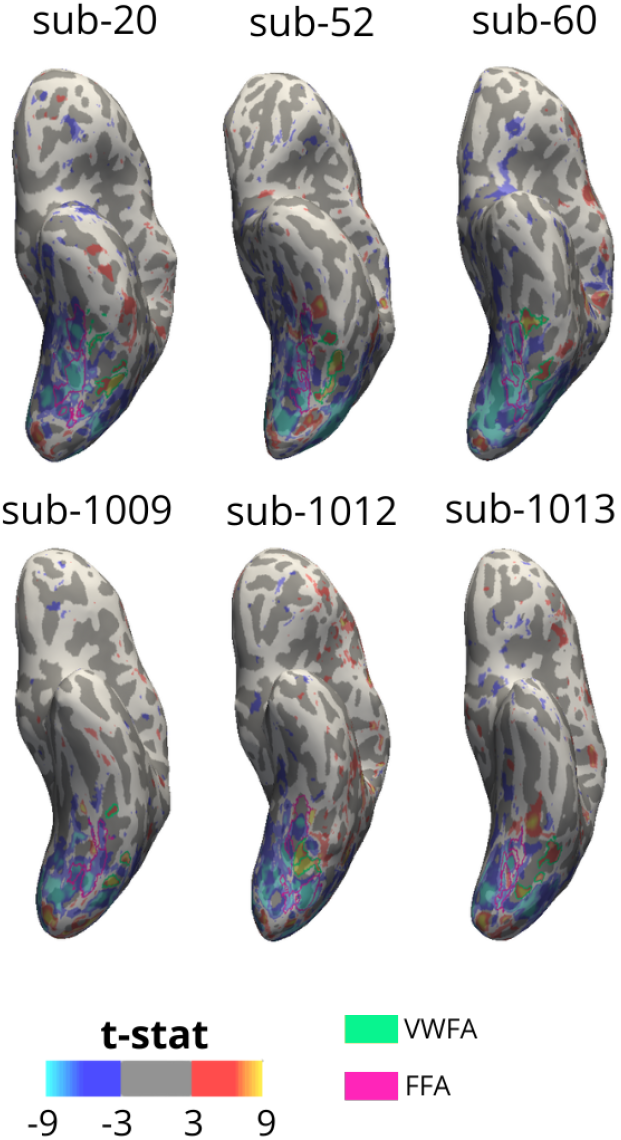
Visual Word Form Area is Smaller and More Variable than Fusiform Face Area. Visual Word Form Areas (VWFAs) and Fusiform Face Areas (FFAs) from three child (top) and three adult (bottom) participants displayed on each participant’s native inflated surface. Heat maps show the results of a vertex-wise t test comparing activation to text (warm tones) relative to all other stimuli (pseudo fonts, faces, objects, limbs; cool tones) at a t > 3. VWFAs (green) were drawn using this contrast on each native surface. FFAs (pink) were drawn using a similar process comparing responses to face to all other stimuli at the same t threshold.

### 3.2 VWFA is Consistent Within Individuals

We next sought to determine if regions identified as the VWFA consistently and stably respond to text across different experiments. To do this, we split our entire dataset in half by task. We defined VWFAs in each individual using, first, only data from the one-back task runs and second, using only data from the fixation task runs. Thus each individual had regions defined on independent data with different task demands. Figure 2a shows the boundaries of these task-specific VWFAs (tVWFAs) in 10 sample participants.

**Figure 2.**
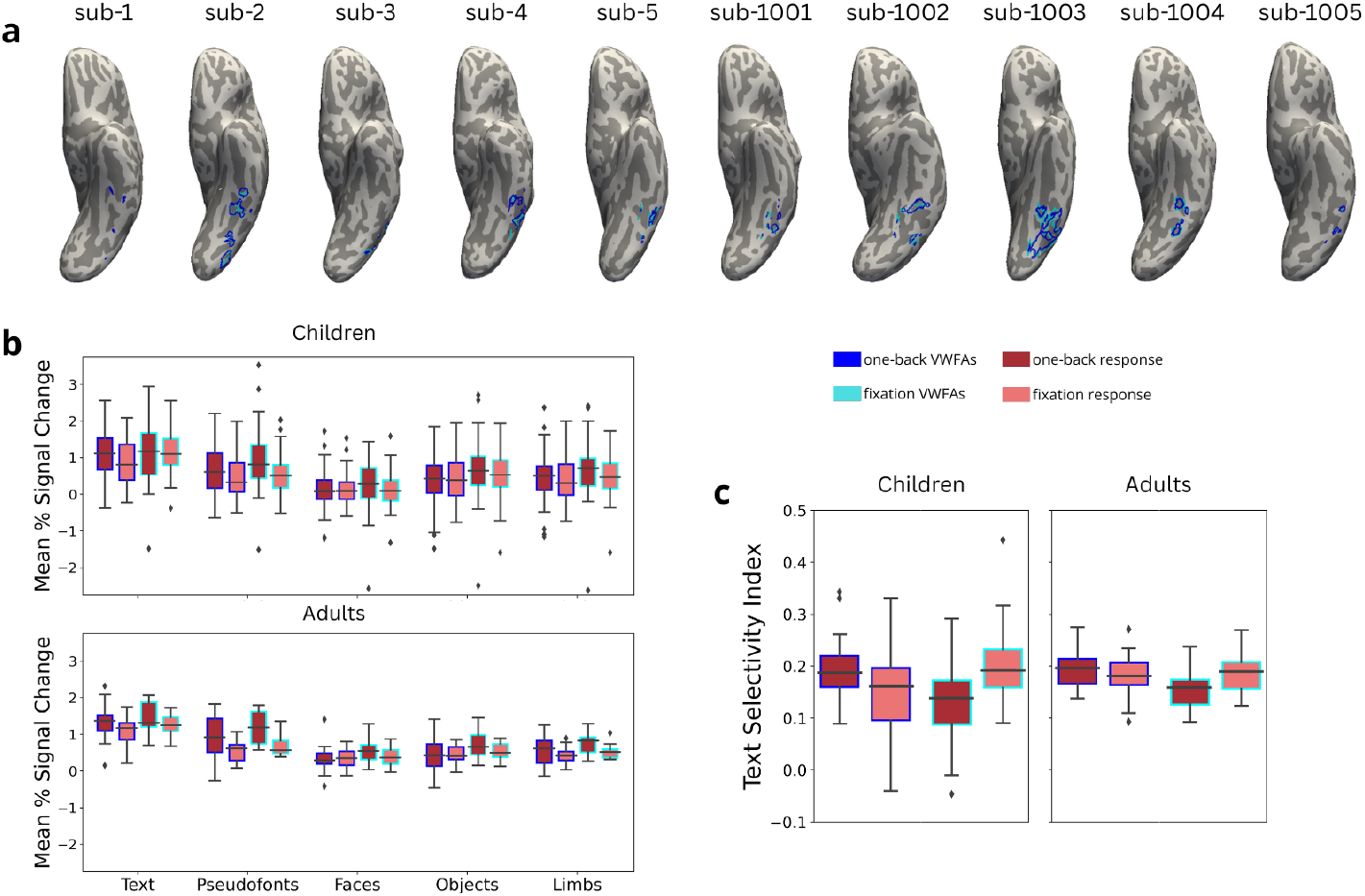
Visual Word Form Area is Selective to Text Regardless of Task Demands. **a**, Visual Word Form Areas (VWFAs) from child (left 5) and adult (right 5) participants displayed on an inflated surface on each participants’ native T1w. Blue VWFAs represent text-selective regions drawn using only data from the one-back task while cyan VWFAs represent text-selective regions drawn using only data from the fixation color change task. **b**, Neural response to visual categories (text, pseudo fonts, faces, objects, and limbs) in units of percent signal change separated by response during each task and within each task-specific VWFA. Data from child participants is displayed in the top plot and data from the adult participants is displayed in the bottom plot. **c**, Neural tuning in terms of a text-selectivity index separated by task and within each task-specific VWFA in children (left) and adults (right). Box plots display the median and quartiles of the data. Outliers are visualized in panels b and c as diamonds above and below the whiskers for each box plot.

To quantify the consistency of the region identified as the VWFA, we calculated a dice similarity coefficient (DSC)^46,47^ to determine how similar one-back tVWFAs (otVWFAs) were to fixation tVWFAs (ftVWFA). We found that children had an average DSC of 0.430 (sd = 0.260, sem = 0.034) and adults had an average DSC of 0.618 (sd = 0.207, sem = 0.057), indicating nearly half of the vertices overlap between tasks within an individual. This suggests that, despite some differences, viewing text consistently activates the same patch of VOTC within the individual (Figure 2a). To contextualize this relationship to the similarity of separate task-evoked regions across categories, we ran an additional analysis calculating the DSC between regions defined for each task separately for each of the other categories in the localizer (faces, objects, limbs, and pseudo fonts). To accomplish this, we took an automated ROI definition approach.

We drew a single VOTC ROI in fsaverage space, projected that label into the native surface of each participant, and grabbed all vertices within that native label that met our previously defined threshold of t > 3 for each respective contrast (i.e. category > all other categories). These results (see Table S1 for more detail) reveal that the DSC values we calculated in the natively defined VWFAs is roughly on par with the those calculated for the other categorically-selective regions (with the exception of a pseudo font region which had very low between-task DSC) defined with an automated approach (i.e. that are not strictly anatomically constrained on native level). We also find that the DSC values are lower in this automated ROI approach when we compare the natively defined VWFA’s to the automated text-selective ROIs, further demonstrating that natively and individually-defined ROIs are more consistent than the automated approach.

We next analyzed whether the ROI defined based on one task was responsive and selective to text across tasks. To accomplish this, we first measured how an individual’s BOLD response to visual stimuli varied between otVWFA and ftVWFA by calculating the percent signal change within an individual for each stimulus category using all available data from both tasks (Figure 2b). A linear mixed effect model revealed, unsurprisingly, a negative main effect for each category in both children (p < 3.11e^-8^) and adults (p < 7.62e^-5^), confirming that response to text was higher than any other category within the reference ROI. Furthermore, the lack of two-way interaction effects between every stimulus category and both ROI task and BOLD task in children (p > 0.056) in children demonstrates that the difference between response to text and response to every other category is consistent during the fixation task and within the ftVWFA. This pattern is mostly replicated in the adult sample (p > 0.155) with the exception of an interaction between the pseudofont category and BOLD task ( β = -0.217, p = 0.025), suggesting that adults may respond to pseudo fonts even less responsive to pseudo fonts during the fixation task. Regardless, adults still demonstrated the pattern of stronger response to text during every combination of ROI task and BOLD task compared to every other category.

By splitting the data in half, we were able to cross validate the text-selectivity of VWFA across tasks. In other words, VWFAs defined with one-back data were still more responsive to text during a fixation task and VWFAs defined with fixation data were also more responsive to text during the one-back task compared to any other stimuli. We can therefore conclude that, even without perfect overlap a text-selective region could be defined regardless of task demands. From this, we can conclude that, despite what some previous research may claim, stimuli (and not task) are the primary driver of activation within VWFA.

### 3.3 Task Modulates VWFA Response

We next examined within-participant differences in tVWFA properties. We first measured the sizes of each tVWFA to determine if the tasks drove differences in the extent of activated cortical territory. tVWFAs defined with one-back task data were slightly larger (children: 185.207 vertices, SD = 274.711; adults: 346.714 vertices, SD = 342.704) than tVWFAs defined with fixation task data (children: 167.756 vertices, SD = 261.382; adults: 336.571 vertices, SD = 417.956). A paired-samples t-test (see Table S2) revealed that this effect was significant in the child sample, (*t*(81) = 2.358, p = 0.021) but not the adult sample (*t*(13) = 0.561, p = 0.584).

Next, we assessed whether the functional tuning properties of tVWFAs (i.e., BOLD response amplitudes within the ROIs) were modulated by task (see Table S3). We found that both groups displayed a significantly positive interaction with the ROI task (children: β = 0.186, p = 0.9.61e^-5^; adults: β = 0.146, p = 0.032) and a significantly negative interaction with the BOLD task, (children: β = -0.161, p = 0.001; adults: β = -0.217, p = 0.002). This suggests that the ftVWFA evokes stronger response to text compared to the otVWFA during the one-back task (the reference BOLD task) and that the fixation task evokes a weaker response to text within the otVWFA (the reference ROI task) compared to the fixation BOLD task. This demonstrates a clear task effect in overall response magnitude to text.

We next explored how tuning properties differed across tasks for each category separately. Effects differed slightly by category for both children and adults (see Table S4), but we found that response to text during the one-back task did not differ between the one-back and fixation tVWFAs in children (β = 0.079, p = 0.221) nor in adults (β = 0.124, p = 0.174). We did however find that within the otVWFA, response to text was weaker during the fixation task when compared to BOLD response during the one-back task in both children (β = -0.257, p = 5.06e^-5^) and adults (β = -0.239, p = 0.011). We also found an interaction effect in children (β = 0.202, p = 0.026) such that response to text was strongest within the ftVWFA during the fixation task (See Figure 2b) but this effect was not observed in our adult sample (p > 0.732).

We then calculated an index to determine text selectivity in the individual by comparing a person’s response to text relative to their response to other categories^29,43^ (Figure 2c). We fit a LME estimating text selectivity as a function of the interaction between tVWFA type and task-specific BOLD response (Table S5). Here, we found that text-selectivity during the one-back task was weaker in the ftVWFA compared to the otVWFA in both children (β = -0.090, p = 4.70e^-5^) and adults (β = -0.041, p = 6.03e^-3^). We also found that within the otVWFA, text selectivity was weaker during the fixation task compared to one-back task in children (β =-0.065, p = 2.08e^-3^), however this was not observed in adults (β = -0.019, p = 0.191). Finally, we found an interaction effect in both children (β = 0.155, p = 7.85e^-7^) and adults (β = 0.051, p = 0.015), which suggests, unsurprisingly, that text-selectivity is highest when looking at the response based on the same data on which the ROI was defined.

Together these results suggest task demand can influence the size and tuning properties of VWFA despite the consistency seen in localization. In other words, we confirm that while this region is consistently responsive to the stimuli regardless of task as shown in the previous section, we also see that VWFA is also modulated by task demand. Therefore, while a text-selective region can be defined consistently, the variability in effect sizes in the literature may arise from differing tasks used to evoke responses within the VWFA.

### 3.4 Group Averages in Template Space Obscure Text-Selective Responses

To determine if a text-selective region of cortex can be analyzed using a group level analysis in template space, we used a conventional approach to conducting statistics on the fsaverage template^41,42,48^. This was done by first creating contrast maps for each individual and for each category and transforming them to the fsaverage template. We then ran a 1-sample t-test to determine the vertices that responded significantly more to each category compared to the others. As seen in Figure 3, the ventral surface displays strong negative t-statistics throughout VOTC in both the text > other and the pseudofonts > other group-level maps, indicating a greater response to other categories and no selectivity for text or text-like stimuli. Meanwhile, VOTC does show highly significant responses for the face > other, objects > other, and limbs > other contrasts. These maps reveal a distinct difference in the way group averaging of activation maps affects the inferences that will be drawn regarding different categories of visual stimuli. Furthermore, these new insights reveal that even though text-selective responses could be localized in each individual (and cross-validated across tasks) these small and anatomically variable regions become obscured in a group average.

**Figure 3.**
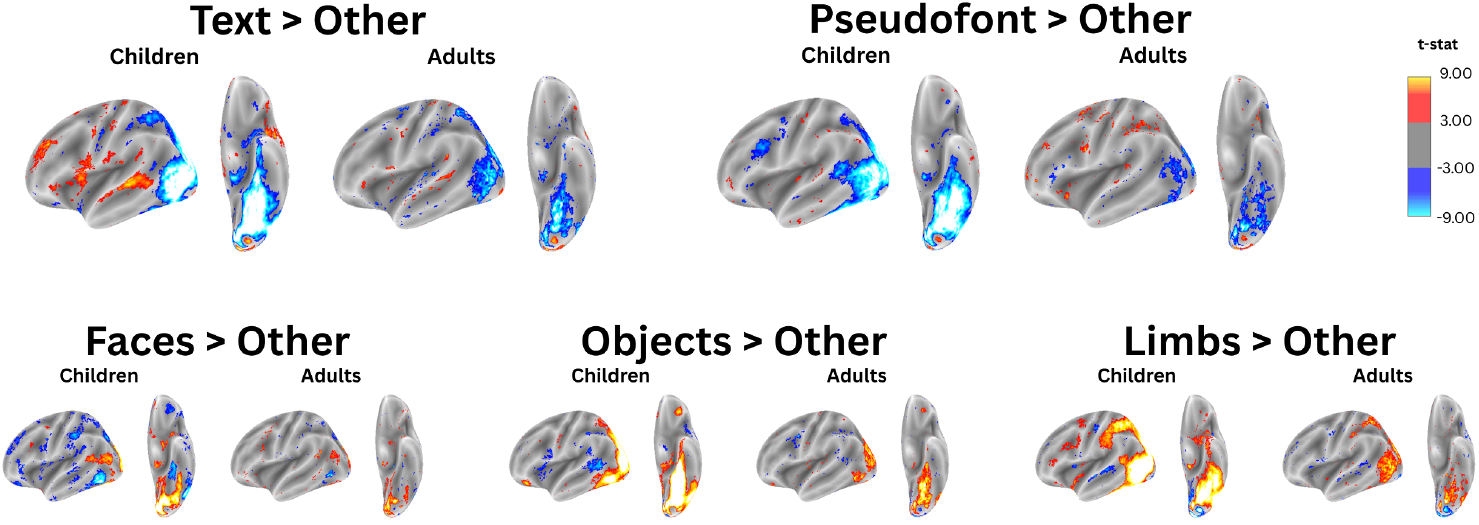
Group Activations Obscure Text-Like Selectivity in High Level Visual Cortex. Group level activation maps in the left hemisphere in children (left) and adults (right) for each contrast category (text, pseudofonts, faces, objects, and limbs). A general linear model was fit to individual participant data to calculate contrast maps for each participant. Group level analysis was completed by performing a one-sample t-test on all contrast maps. Maps are thresholded at a t of +/- 3 with all positive values displaying selectivity for the named stimulus category (warm tones) and all negative values displaying selectivity of all other, unnamed stimulus categories (cool tones).

### 3.5 Group and Template VWFAs Obscure Text-Selective Results Due to Individual Differences

Next, we asked whether group-level ROIs, defined on a template, could capture the same response properties as natively drawn individual ROIs. In line with other studies that have defined and published template ROIs^44^, we first created group average ROIs for both VWFA and FFA for children and adults. To this end, each participant’s VWFA and FFA were projected to fsaverage (see Figure 4a). We then calculated a probability map representing the percentage of participants with an overlapping ROI at every fsaverage vertex in the left hemisphere. These heat maps, seen in Figure 4b, represent the areas of cortex a given ROI was found in any participant. These maps show that there was not a single vertex that contained the VWFA or FFA in all participants (100% probability). Thus, both FFA and VWFA vary significantly across participants - a result that mirrors previous findings^49^. However, fewer participants shared VWFA vertices (maximum of 25.7% in children and 67.1% in adults) than FFA vertices (maximum of 46.2% in children and 78.6% in adults).

**Figure 4.**
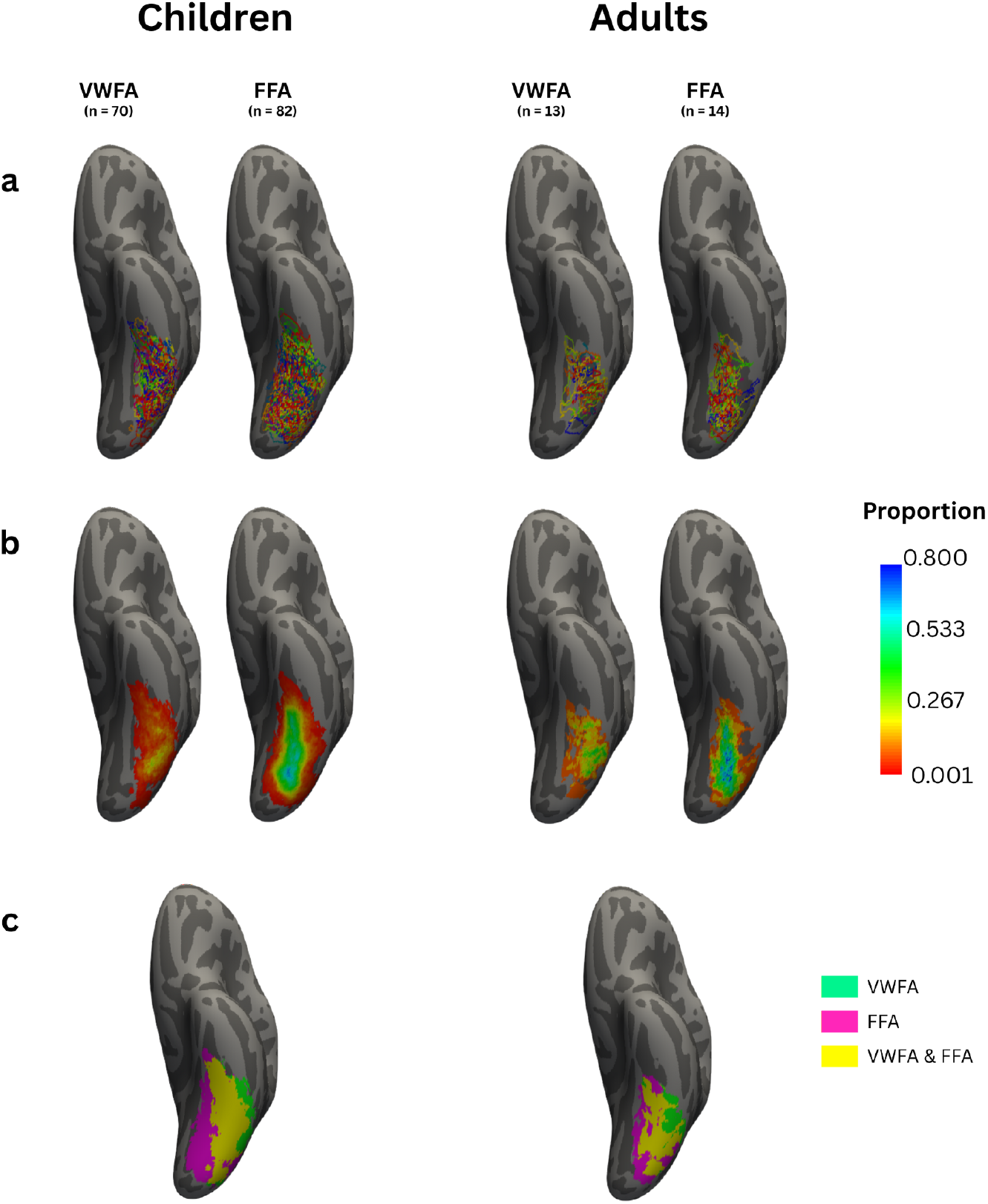
Variability in Text and Face Selective Regions Across Individuals. **a**, native Visual Word Form Area labels (VWFA; left) and Fusiform Face Area labels (FFA; right) projected into average space and projected onto an average surface template for every participant in children (left) and adults (right). Different colors represent a label from a different participant. **b**, Heat maps displaying the proportion of participants who had a given region at every vertex. **c**, Indicator map displaying vertices where a VWFA (green) or an FFA (pink) was present in at least one participant in each group. Any vertex in which at least 1 participant had a VWFA and at least 1 other participant had a FFA is displayed in yellow.

We next examined the variability in the spatial configuration of these regions. Figure 4c visualizes the overlap of text and face ROIs across all participants. This analysis revealed that the area of cortex that is text-selective in some participants, while also being face-selective in others, far outweighs the amount of cortex that is uniquely selective to each category separately. Moreover, the region of cortex that is only responsive to text is substantially smaller than the region of cortex that is only responsive to faces. The amount of cortex that is strictly text-selective is equivalent to 649 vertices in children and 962 vertices in adults (equivalent to about 0.4% and 0.8% of the total 163,842 vertices in the fsaverage template), compared to the 1,369 vertices in children and 2,450 in adults (0.8% and 1.5%) that are strictly face-selective.

Finally, 2,026 vertices in adults and 3,093 vertices in children (1.2% and 1.9%) belong to cortical territory that is text-selective in some participants but face-selective in others. Thus, nearly twice as many vertices are strictly face selective than are strictly text selective.

To further investigate the extent of variability in the topography of the VWFA, we performed a series of analyses to investigate within-participant size differences and between participant differences in ROI center. We first calculate the size for each participant’s manually defined VWFA and FFA. Using a one-sample t-test, we find that VWFA is significantly smaller than the FFA in both children (t(81) = -11.685, p = 4.65e^-19^) and adults (t(13) = -2.614, p = 0.021; see Table S6). We next sought to determine if there was more spatial variability in the VWFA location across participants compared to the spatial variability of the FFA. To do this, we first projected each participant’s native label to the fsaverage surface. We then calculated the medoid (ROI center) of each participant’s ROI by identifying the within-ROI vertex that minimized the distance to all other vertices within the ROI. Once the ROI center was defined for each participant, we then calculated a group medoid by using the same process to identify the medoid vertex that minimized the distances to all other participant medoids. We find that the average distance from any participants VWFA medoid to the group VWFA medoid is 13.904 mm (SD = 10.024) in children and 10.9125 mm (SD = 10.024) in adults. Additionally, the average distance from any participants FFA to the group FFA medoid is 10.191 mm (SD = 8.595) in children and 9.040 mm (SD = 6.754) in adults. Furthermore, a two-sample t-test reveals that in the child sample, the participant medoid to group medoid distance is larger for VWFAs compared to FFAs (t(162) = 4.714, p = 5.20e^-6^; see Table S7). However, this effect was present but not significant in the adult sample ( t(26) = 0.830, p = 0.414). Combined, these results indicate that on average, VWFA is both smaller within individuals and more variable in spatial location across individuals compared to the immediately adjacent control region (FFA).

Despite this large amount of between-participant VWFA-FFA overlap and the evidence of heightened variability in VWFA size and location, we followed the standard approach to create group-average ROIs in order to determine if group-derived labels can accurately measure text-selective responses when applied to individual level activation maps. Specifically, ROIs were created by taking all vertices from the probability map that displayed at least 20% of participant overlap^43,44^. We created separate labels for child and adult participants in case any developmental differences exist between the two age groups. We used template ROIs from these recent studies to compare results across several examples of group-level ROI analytical choices. We then analyzed the response properties of these different ROIs (see Figure 5a) to elucidate how these methodological choices affect the inferences that are made about the reading circuitry.

**Figure 5.**
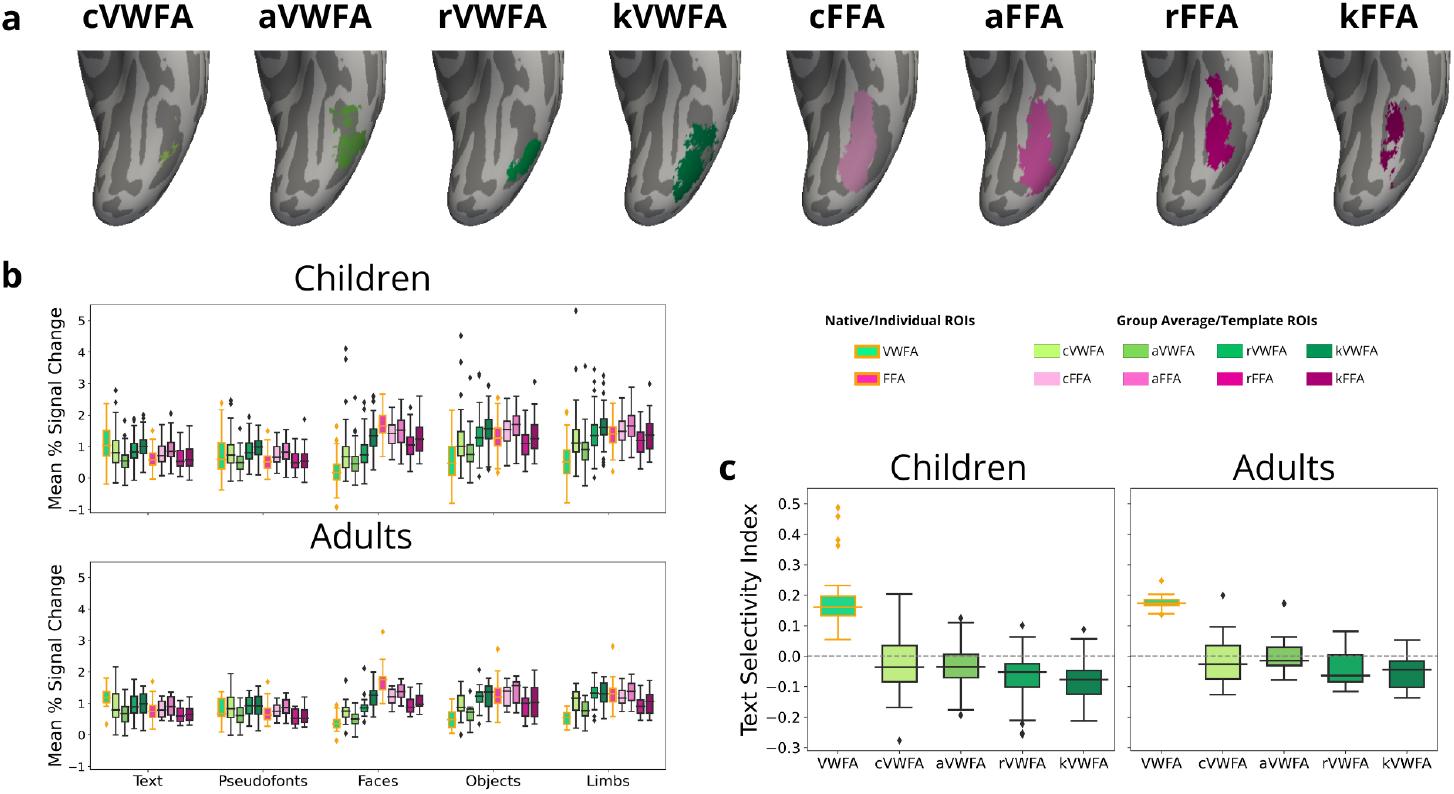
Group Template ROIs Fail to Capture Individual-Level Text Response. **a** Each of the four group average and template Visual Word Form Areas (VWFA) and Fusiform Face Areas (FFA) displayed on an inflated ventral surface of the fsaverage anatomy. **b** Response profiles for each of the average and template VWFAs and FFAs displayed next to the response profile based on individually-defined native VWFAs and FFAs (highlighted in orange). Response is measured in terms of percent signal change from a blank condition. Data from child participants is displayed in the top plot and data from the adult participants is displayed in the bottom plot. **c** Tuning properties of each of the group and template VWFAs calculated as a text-selectivity index displayed with the text selectivity index extracted from individually-defined native VWFAs.Data from child participants is displayed in the left plot and data from the adult participants is displayed in the right plot. VWFA = values extracted from natively defined VWFA; FFA = values extracted from natively defined FFA; c = group average ROI of study child participants; a = group average ROI of study adult participants; r = template ROIs from Rosenke et al., 2018; k = template ROIs from Kubota et al., 2023

We first wanted to determine if the amplitude of the response to words is lower and the response to other categories was higher in group and template ROIs compared to native ROIs (Figure 5b). We found that most group and template VWFAs measured weaker responses to text than the native VWFA in both children (p < 0.005) and adults (p = 0.031) with the exception of the Kubota et al. VWFA (kVWFA) which showed no difference in text response compared with the native VWFA (children: p = 0.841; adults: p = 0.104; see Table S8). This confirmed our hypothesis that group and template ROIs won’t capture peak mean response to text. For pseudofonts, we found mixed results: Compared with the native VWFA, the child group VWFA (cVWFA) showed no difference in response within the child sample (β = 0.089, p = 0.096) and the adult sample (β = 0.072, p = 0.433), the adult group VWFA (aVWFA) showed a lower response in both children (β = -0.179, p = 0.001) and adults (β = -0.188, p = 0.046), and both the Rosenke et al. VWFA (rVWFA) and kVWFA showed a higher response in children (rVWFA: β = 0.161, p = 0.003, kVWFA: β = 0.280, p = 2.80e^-7^) but not in adults (rVWFA: β = 0.139, p = 0.134, kVWFA: β = 0.157, p = 0.092). As expected, VWFAs showed greater response magnitude to faces, objects, and limbs in all group/template ROIs in children (p < 0.001; see Table S8a) and in most group/template ROIs in adults (p < 0.001; see Table S8b). This is in line with the notion that text-responses get washed out when averaged across participants.

Combined, this shows that response magnitude to every stimulus category varies greatly depending on the ROI used to analyze this form of functional tuning (see Tables S9a and S9b for more information on FFAs).

We then repeated the text-selectivity index calculation from earlier to determine if the group and template VWFAs accurately identified a patch of cortex that was indeed text selective in our participants. Figure 5c shows that while the individually drawn native ROIs are indeed text-selective (i.e. their selectivity indices are positive as expected given they were defined using a text-selective contrast map), all four of our group and literature-based template ROIs failed to capture text-selective cortex in our participants, showing a negative median selectivity index. Fitting a LME to this data to quantify this difference, we found a negative effect for each group and template VWFA in both children (p < 2.09e^-47^) and adults (p < 5.16e^-12^), confirming with a high degree of significance that none of these group/template VWFAs were text selective.

The combination of the BOLD response analysis and the selectivity index analysis explains the lack of a text-selective response seen in Figure 3. When one patch of cortex is selective to one category in some individuals but is also selective to a different category in others, averaging responses across individuals will “wash out” category-specific responses. This comprehensive approach offers further evidence to support existing literature which suggests that group-level approaches are inadequate for analyzing text-selective responses in VOTC.

## 4 Discussion

This study investigated how methodological choices affect the inferences that are drawn about text-selective responses in VOTC. We first used a functional localizer to confirm that we could localize the VWFA within the mosaic of category selective regions in VOTC. We manually defined ROIs for each category-selective region on the native cortical surface and confirmed that, within each individual’s brain, a consistent patch of cortex selectively responded to words irrespective of the task. While we found evidence that the task demands affected the size, response magnitude, and selectivity within the region, we also confirmed that VWFA was consistently localized within an individual across. We therefore conclude that the presence of text stimuli, and not the task demands as previous research may suggest^11^, primarily drive the response within VWFA.

We then investigated how moving from an analysis of individual brains to an analysis in template space (fsaverage cortical surface) affected the inferences we might draw about text-selective responses in VOTC. We found that while faces, objects, and limbs evoked a response in a consistent region of cortex across participants, a text-selective cortex could not be identified in a group average despite its existence on an individual level. This suggests that anatomical and functional topography varies significantly across individuals. To understand this relationship better, we looked at how individual ROIs overlap and found that VWFAs across individuals vary in anatomical location. We found that patches of cortex that respond to text in some individuals respond to different visual categories in other individuals. This was even more pronounced in children. In a group level analysis, these conflicting ROIs that share cortical territory will neutralize text-selective responses when averaging across individuals. Additionally, we showed the importance of an individual approach to studying neural response to text by exploring nullifying results obtained from a group level ROI definition. From these findings, we conclude that, with the unique nature of smaller and more variable text-selective regions, individual approaches to analysis in the native space are crucial to obtain consistent and reproducible results. These conclusions are in line with Glezer and Riesenhuber^26^), further emphasizing the importance of an individual analysis approach and encouraging future researchers to follow this practice. However, it is important to note that the process of group registration and normalization itself may introduce inaccuracies in the functional and anatomical alignment, which could contribute to the blurring of results in group analyses in addition to individual differences.

Previous research suggests that certain functionally-selective cortical patches for categories such as faces, limbs, and places in the VOTC are present even without access to visual experience^50^, and that this organization is the result of basic needs and the ecologically advantageous nature of efficient object discrimination^51^. As children grow, changes in their categorically-selective ROIs suggest a recycling model in which selectivity for text increases while selectivity for another category may decrease^36,52,53^. This cortical recycling theory suggests that there is competition for cortical territory in the VOTC which could lead to the variability in VWFA localization observed in this study. Text recognition is a learned task that takes instruction and practice over years of development, and has only become common practice in the very recent evolutionary timeline compared to other categories of object recognition.

Individual-specific experiences may shape the way this patch of cortex develops more than other known categorically selective regions because of its delayed emergence compared to other studied regions. Further research is needed to understand how the VWFA emerges as people learn how to read, and the effect this development has on the VOTC and other contributing brain regions. Longitudinal studies allow us to measure the changes/emergence of VWFA in individuals as children grow up and gain reading skills. We recently showed that, using individually defined VWFA, we were able to capture growth in VWFA related to growth in reading ability and found a strong relationship between VWFA size and reading ability^29^.

We argue that an accurate and comprehensive understanding of VWFA function necessitates its definition individually on native anatomy. Though this argument has been made previously^26^, we have yet to see consistent adherence to this guidance. Here we provide a deeper dive into sources of consistency within individuals and variability across individuals.

VWFA exhibits consistent localization within individuals despite differences that may arise due to specific task demands utilized to selectively activate the region. Furthermore, group-level functional data and template-based ROIs are inherently inadequate for precisely identifying text-selective cortex within the ventral occipitotemporal cortex. This is particularly pronounced because text represents a unique case of functional specialization compared to other visual categories, such as faces, which have a longer evolutionary history of recognition. Our results suggest that a transition from template-based analyses to individually-localizing regions of interest will resolve ambiguities and controversies that persist in the literature. Particularly in the study of development, an individual approach will be critical for understanding the mechanisms of stability and change within high-level visual cortex.

## 5 Data and Code Availability

De-identified data has been made publicly available through the Stanford University Libraries Digital Repository and can be found at https://doi.org/10.25740/jp789mc0395. Code has been made publicly available through an online GitHub repository and can be found at https://github.com/jamielmitchell/Mitchell_VWFAmethods_2025.git.

## 6 Author Contributions

**JLM:** Conceptualization, Formal analysis, Investigation, Writing (original draft & review and editing), Visualization. **MFJ:** Investigation, Data curation, Project administration. **HLS:** Investigation, Data curation, Project administration. **MY:** Investigation, Supervision. **JDY:** Conceptualization, Resources, Writing (review and editing), Supervision, Funding acquisition.

## 9 Acknowledgements

We would like to thank the individuals and families who participated in this study and the research coordinators and assistants who contributed to data collection.

