## Supplemental Material for "Visual Word Form Area demonstrates individual and task-agnostic consistency but inter-individual variability"

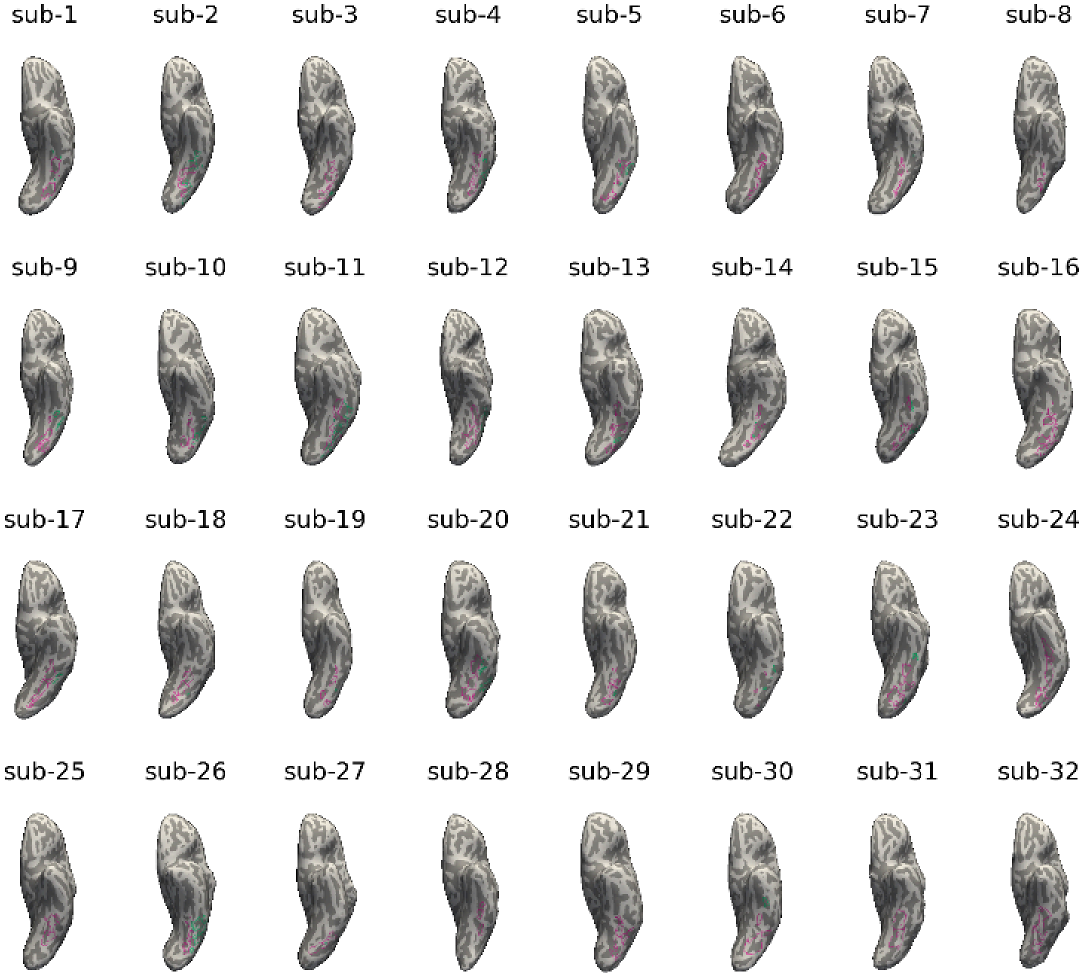


**Figure S1 | Individual Participant Regions of Interest - Subs 1-32**
Visual Word Form Area (VWFA; green) and Fusiform Face Area (FFA; pink) displayed on a ventral view of the inflated surface of the left hemisphere for every participant in the study.

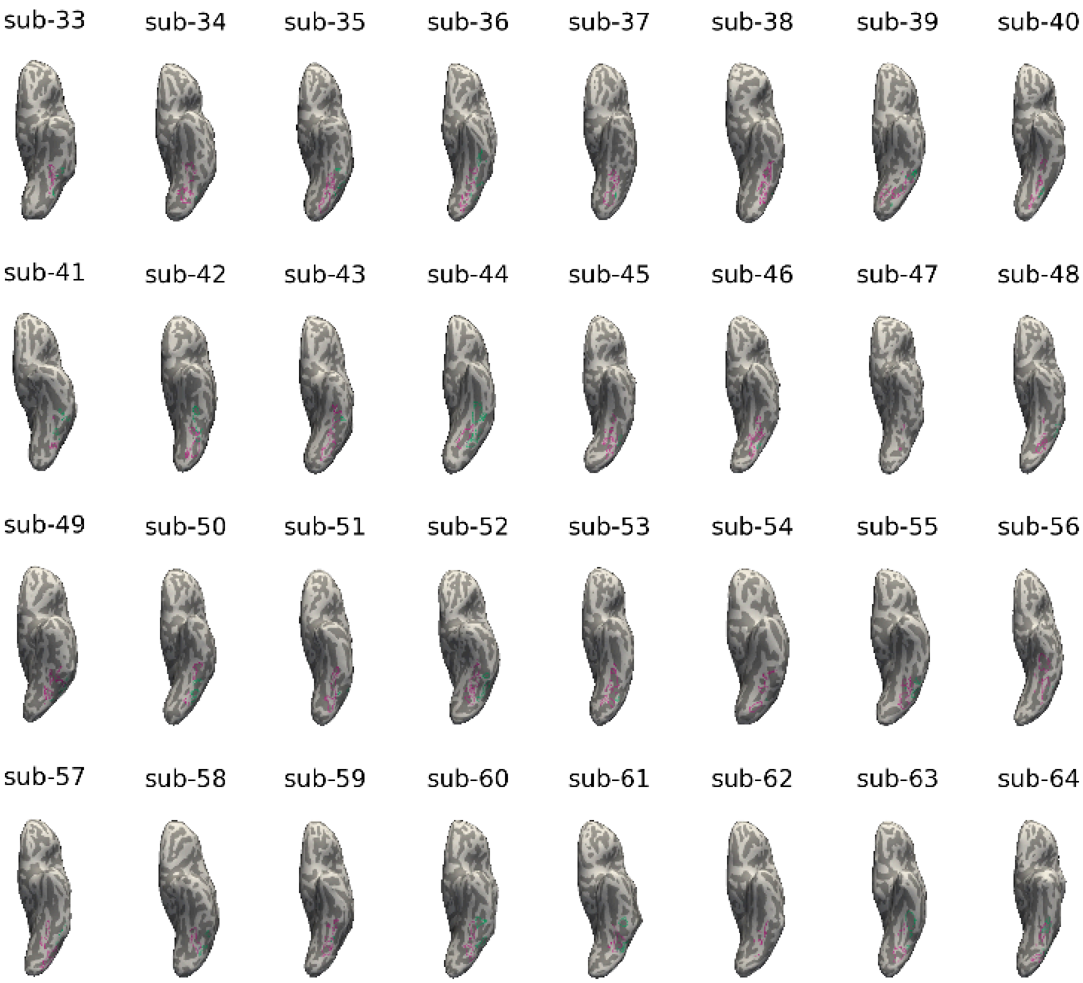


**Figure S2 | Individual Participant Regions of Interest - Subs 3-64**
Visual Word Form Area (VWFA; green) and Fusiform Face Area (FFA; pink) displayed on a ventral view of the inflated surface of the left hemisphere for every participant in the study.

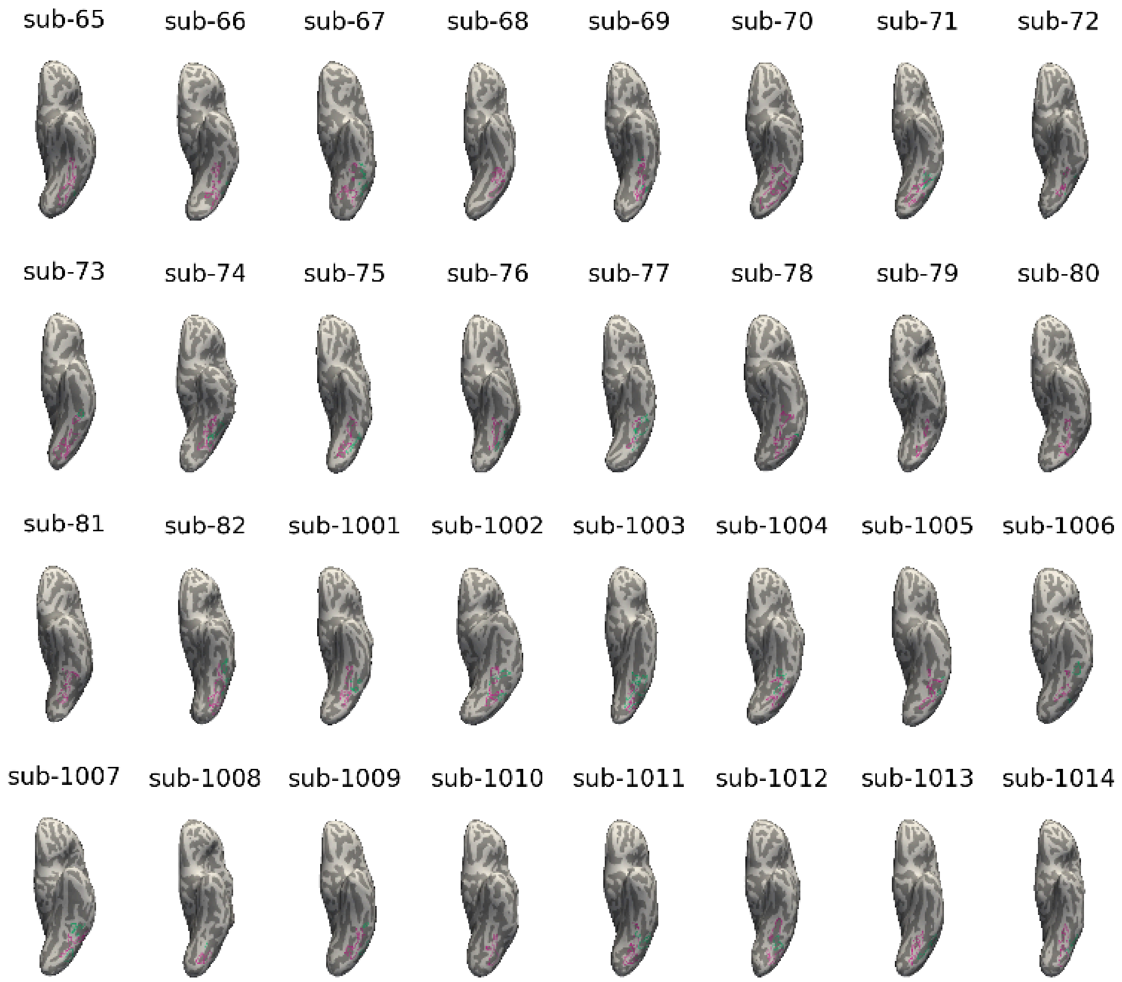


**Figure S3 | Individual Participant Regions of Interest - Subs 65-82 & 1000-1014**
Visual Word Form Area (VWFA; green) and Fusiform Face Area (FFA; pink) displayed on a ventral view of the inflated surface of the left hemisphere for every participant in the study.

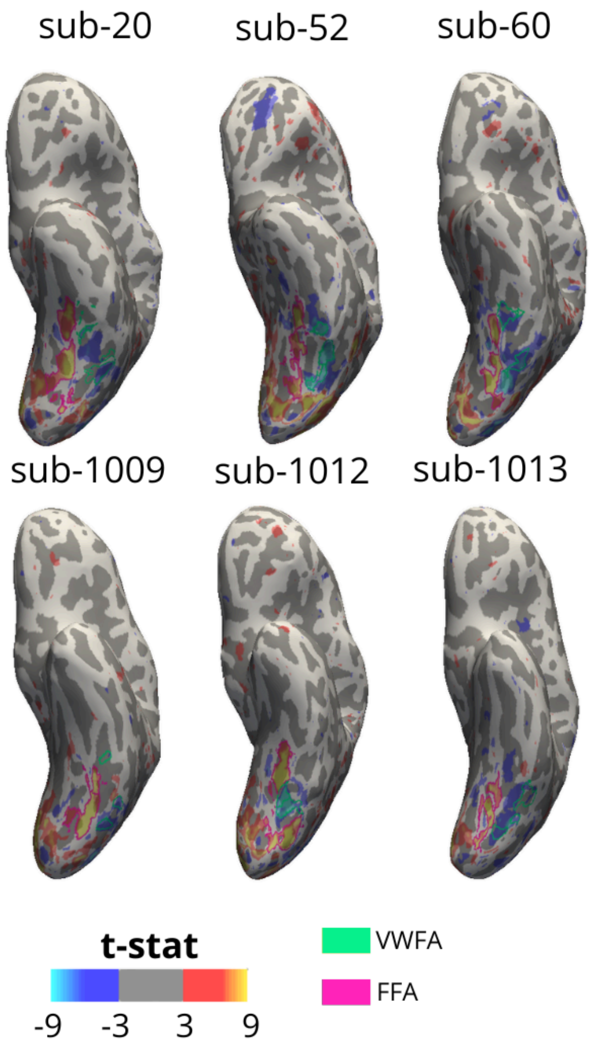


**Figure S4 | Visual Word Form Area is Smaller and More Variable than Fusiform Face Area (Face Maps)**
Visual Word Form Areas (VWFAs) and Fusiform Face Areas (FFAs) from three child (top) and three adult (bottom) participants displayed on each participant's native inflated surface. Heat maps show the results of a vertex-wise t test comparing activation to faces (warm tones) relative to all other stimuli (text, pseudo fonts, objects, limbs; cool tones) at a t > 3. VWFAs (green) were drawn using this contrast on each native surface. FFAs (pink) were drawn using a similar process comparing responses to face to all other stimuli at the same t threshold.

| SepTask VWFA Dice Similarity Coefficient | | | |
|---|---|---|---|
| | **Mean DSC** | **SD** | **SEM** |
| **VWFA (native)** | | | |
| Children | 0.430 | 0.260 | 0.034 |
| Adults | 0.618 | 0.207 | 0.057 |
| **Text Area** | | | |
| Children | 0.336 | 0.271 | 0.031 |
| Adults | 0.558 | 0.236 | 0.063 |
| **Faces Area** | | | |
| Children | 0.579 | 0.191 | 0.021 |
| Adults | 0.657 | 0.118 | 0.031 |
| **Objects Area** | | | |
| Children | 0.674 | 0.184 | 0.020 |
| Adults | 0.696 | 0.104 | 0.028 |
| **Limbs Area** | | | |
| Children | 0.533 | 0.190 | 0.021 |
| Adults | 0.460 | 0.150 | 0.040 |
| **Pseudo Fonts Area** | | | |
| Children | 0.078 | 0.157 | 0.022 |
| Adults | 0.020 | 0.027 | 0.008 |

**Table S1 | Dice Similarity Calculations for Split Task Regions of Interest**
Results from a Dice Similarity calculation for computing the overlap of regions of interest (ROIs) when drawn using only one-back task data and only fixation task data. Contrast maps were created for each category of stimuli with a category > all other categories contrast and thresholded at t > 3. VWFAs were defined manually for each participant and included all vertices greater than threshold with strict native anatomical boundaries. Test, Faces, Objects, Limbs, and Pseudo Fonts Areas were defined with an automated approach grabbing all vertices greater than threshold within an average ventral occipitotemporal cortex label boundary.

| SepTask VWFA Size Difference | | | | | |
|---|---|---|---|---|---|
| | **t** | **CI** | | **DOF** | ***p*** |
| | *(ob > fix)* | *low* | *high* | | |
| Children | **2.358** | **0.029** | **0.339** | **81** | **0.021** * |
| Adults | 0.561 | -0.127 | 0.216 | 13 | 0.584 |

**Table S2 | Task Differences in Visual Word Form Area Size**
Results from a two-tailed, paired-samples t-test comparing the size of the VWFA (in $\log_{10}$ number of vertices) when drawn using only one-back task data and only fixation task data. Significant results are displayed in bold and asterisks indicate the degree of significance ($p < 0.001$: ***, $p < 0.01$: **, $p < 0.05$: *)

**a**

| BOLD Response ~ Category * ROI Task + Category * BOLD Task + (1 \| Participant) - Children | | | | | |
|---|---|---|---|---|---|
| | β | Std. Err | DOF | t | p |
| Intercept (ROI Task: One-back \| Map Task: One-back \| Text) | **1.050** | **0.074** | **123.598** | **14.150** | **1.28E-27*** ** |
| Category: Pseudofonts | **-0.317** | **0.057** | **1158.749** | **-5.573** | **3.11E-08*** ** |
| Category: Faces | **-0.856** | **0.057** | **1158.749** | **-15.031** | **8.75E-47*** ** |
| Category: Objects | **-0.598** | **0.057** | **1158.749** | **-10.505** | **1.01E-24*** ** |
| Category: Limbs | **-0.522** | **0.057** | **1158.749** | **-9.167** | **2.15E-19*** ** |
| ROI Task: Fixation | **0.186** | **0.048** | **1160.758** | **3.914** | **9.61E-05*** ** |
| Map Task: Fixation | **-0.161** | **0.047** | **1158.749** | **-3.418** | **0.001*** ** |
| Category: Pseudofonts * ROI Task: Fixation | -0.021 | 0.067 | 1158.749 | -0.312 | 0.755 |
| Category: Faces * ROI Task: Fixation | -0.128 | 0.067 | 1158.749 | -1.916 | 0.056 |
| Category: Objects * ROI Task: Fixation | 0.011 | 0.067 | 1158.749 | 0.160 | 0.873 |
| Category: Limbs * ROI Task: Fixation | -0.012 | 0.067 | 1158.749 | -0.178 | 0.859 |
| Category: Pseudofonts * Map Task: Fixation | -0.106 | 0.067 | 1158.749 | -1.593 | 0.111 |
| Category: Faces * ROI Map: Fixation | 0.078 | 0.067 | 1158.749 | 1.168 | 0.243 |
| Category: Objects * ROI Map: Fixation | 0.096 | 0.067 | 1158.749 | 1.437 | 0.151 |
| Category: Limbs * ROI Map: Fixation | 0.014 | 0.067 | 1158.749 | 0.203 | 0.839 |

**b**

| BOLD Response ~ Category * ROI Task + Category * BOLD Task + (1 \| Participant) - Adults | | | | | |
|---|---|---|---|---|---|
| | β | Std. Err | DOF | t | p |
| Intercept (ROI Task: One-back \| Map Task: One-back \| Text) | **1.318** | **0.101** | **25.161** | **13.001** | **1.15E-12*** ** |
| Category: Pseudofonts | **-0.334** | **0.083** | **233.000** | **-4.028** | **7.62E-05*** ** |
| Category: Faces | **-0.931** | **0.083** | **233.000** | **-11.206** | **1.35E-23*** ** |
| Category: Objects | **-0.839** | **0.083** | **233.000** | **-10.107** | **3.78E-20*** ** |
| Category: Limbs | **-0.727** | **0.083** | **233.000** | **-8.760** | **4.09E-16*** ** |
| ROI Task: Fixation | **0.146** | **0.068** | **233.000** | **2.153** | **0.032*** |
| Map Task: Fixation | **-0.217** | **0.068** | **233.000** | **-3.196** | **0.002** ** |
| Category: Pseudofonts * ROI Task: Fixation | 0.012 | 0.096 | 233.000 | 0.121 | 0.904 |
| Category: Faces * ROI Task: Fixation | -0.017 | 0.096 | 233.000 | -0.176 | 0.860 |
| Category: Objects * ROI Task: Fixation | 0.030 | 0.096 | 233.000 | 0.314 | 0.754 |
| Category: Limbs * ROI Task: Fixation | 0.003 | 0.096 | 233.000 | 0.031 | 0.975 |
| Category: Pseudofonts * Map Task: Fixation | **-0.217** | **0.096** | **233.000** | **-2.263** | **0.025*** |
| Category: Faces * ROI Map: Fixation | 0.137 | 0.096 | 233.000 | 1.427 | 0.155 |
| Category: Objects * ROI Map: Fixation | 0.131 | 0.096 | 233.000 | 1.362 | 0.174 |
| Category: Limbs * ROI Map: Fixation | 0.024 | 0.096 | 233.000 | 0.252 | 0.801 |

**Table S3 | LME Results for Activation by Split Task VWFA on Split Task Data**
Linear mixed effects model results for model predicting BOLD response as a function of the interaction between stimulus category and task-specific ROI along with the interaction between stimulus category and task-specific BOLD. Results are displayed in child participants (**a**) and adult participants (**b**) separately. Significant results are displayed in bold and asterisks indicate the degree of significance ($p < 0.001$: ***, $p < 0.01$: **, $p < 0.05$: *)

**a**

| BOLD Response ~ ROI Task * Map Task + (1 \| Participant) - Children | β | Std. Err | DOF | t | p |
|---|---|---|---|---|---|
| **Text** | | | | | |
| Intercept (ROI Task: One-back \| Map Task: One-back) | **1.094** | **0.080** | **105.338** | **13.724** | **3.39E-25*** ** |
| ROI Task: Fixation | 0.079 | 0.065 | 180.107 | 1.229 | 0.221 |
| Map Task: Fixation | **-0.257** | **0.062** | **177.646** | **-4.154** | **5.06E-05*** ** |
| ROI Task: Fixation * Map Task: Fixation | **0.202** | **0.090** | **177.646** | **2.247** | **0.026*** |
| **Pseudofonts** | | | | | |
| Intercept (ROI Task: One-back \| Map Task: One-back) | **0.694** | **0.080** | **111.418** | **8.645** | **4.49E-14*** ** |
| ROI Task: Fixation | **0.251** | **0.069** | **179.481** | **3.617** | **3.87E-04*** ** |
| Map Task: Fixation | **-0.193** | **0.067** | **176.629** | **-2.900** | **0.004** ** |
| ROI Task: Fixation * Map Task: Fixation | -0.157 | 0.096 | 176.629 | -1.630 | 0.105 |
| **Faces** | | | | | |
| Intercept (ROI Task: One-back \| Map Task: One-back) | **0.156** | **0.064** | **114.972** | **2.426** | **0.017*** |
| ROI Task: Fixation | **0.141** | **0.057** | **180.258** | **2.477** | **0.014*** |
| Map Task: Fixation | -0.015 | 0.054 | 177.294 | -0.284 | 0.777 |
| ROI Task: Fixation * Map Task: Fixation | -0.143 | 0.079 | 177.294 | -1.808 | 0.072 |
| **Objects** | | | | | |
| Intercept (ROI Task: One-back \| Map Task: One-back) | **0.424** | **0.081** | **108.712** | **5.217** | **8.75E-07*** ** |
| ROI Task: Fixation | **0.249** | **0.068** | **180.254** | **3.656** | **3.36E-04*** ** |
| Map Task: Fixation | -0.002 | 0.065 | 177.622 | -0.033 | 0.974 |
| ROI Task: Fixation * Map Task: Fixation | -0.133 | 0.095 | 177.622 | -1.407 | 0.161 |
| **Limbs** | | | | | |
| Intercept (ROI Task: One-back \| Map Task: One-back) | **0.497** | **0.080** | **104.055** | **6.244** | **9.40E-09*** ** |
| ROI Task: Fixation | **0.244** | **0.064** | **179.563** | **3.808** | **1.92E-04*** ** |
| Map Task: Fixation | -0.083 | 0.061 | 177.123 | -1.354 | 0.177 |
| ROI Task: Fixation * Map Task: Fixation | -0.136 | 0.089 | 177.123 | -1.521 | 0.130 |

**b**

| BOLD Response ~ ROI Task * Map Task + (1 \| Participant) - Adults | β | Std. Err | DOF | t | p |
|---|---|---|---|---|---|
| **Text** | | | | | |
| Intercept (ROI Task: One-back \| Map Task: One-back) | **1.329** | **0.123** | **18.294** | **10.803** | **2.26E-09*** ** |
| ROI Task: Fixation | 0.124 | 0.089 | 36.000 | 1.388 | 0.174 |
| Map Task: Fixation | **-0.239** | **0.089** | **36.000** | **-2.667** | **0.011*** |
| ROI Task: Fixation * Map Task: Fixation | 0.044 | 0.126 | 36.000 | 0.345 | 0.732 |
| **Pseudofonts** | | | | | |
| Intercept (ROI Task: One-back \| Map Task: One-back) | **0.959** | **0.120** | **18.947** | **7.964** | **1.83E-07*** ** |
| ROI Task: Fixation | **0.207** | **0.091** | **36.000** | **2.281** | **0.029*** |
| Map Task: Fixation | **-0.384** | **0.091** | **36.000** | **-4.224** | **1.56E-04*** ** |
| ROI Task: Fixation * Map Task: Fixation | -0.099 | 0.129 | 36.000 | -0.773 | 0.445 |
| **Faces** | | | | | |
| Intercept (ROI Task: One-back \| Map Task: One-back) | **0.358** | **0.095** | **19.220** | **3.756** | **0.001**** |
| ROI Task: Fixation | **0.188** | **0.073** | **36.000** | **2.582** | **0.014*** |
| Map Task: Fixation | -0.021 | 0.073 | 36.000 | -0.282 | 0.780 |
| ROI Task: Fixation * Map Task: Fixation | -0.119 | 0.103 | 36.000 | -1.150 | 0.258 |
| **Objects** | | | | | |
| Intercept (ROI Task: One-back \| Map Task: One-back) | **0.446** | **0.103** | **21.855** | **4.314** | **2.83E-04*** ** |
| ROI Task: Fixation | **0.242** | **0.089** | **36.000** | **2.725** | **0.010**** |
| Map Task: Fixation | -0.020 | 0.089 | 36.000 | -0.230 | 0.820 |
| ROI Task: Fixation * Map Task: Fixation | -0.131 | 0.125 | 36.000 | -1.047 | 0.302 |
| **Limbs** | | | | | |
| Intercept (ROI Task: One-back \| Map Task: One-back) | **0.567** | **0.089** | **23.089** | **6.355** | **1.71E-06*** ** |
| ROI Task: Fixation | **0.196** | **0.080** | **36.000** | **2.456** | **0.019*** |
| Map Task: Fixation | -0.145 | 0.080 | 36.000 | -1.822 | 0.077 |
| ROI Task: Fixation * Map Task: Fixation | -0.094 | 0.113 | 36.000 | -0.834 | 0.410 |

**Table S4 | Category-Specific LME Results for Activation by Split Task VWFA on Split Task Data**

Linear mixed effects model results for BOLD response prediction as a function of the interaction between task-specific ROI and task-specific BOLD for each category of experiment stimulus. Results are displayed in child participants (**a**) and adult participants (**b**) separately. Significant results are displayed in bold and asterisks indicate the degree of significance ($p < 0.001$: ***, $p < 0.01$: **, $p < 0.05$: *)

| Text Selectivity ~ ROI Task * Map Task + (1 \| Participant) - Children | | | | | |
|---|---|---|---|---|---|
| | β | Std. Err | DOF | t | p |
| **Children** | | | | | |
| Intercept (ROI Task: One-back \| Map Task: One-back) | **0.215** | **0.016** | **237.509** | **13.754** | **4.70E-32*** ** |
| ROI Task: Fixation | **-0.090** | **0.022** | **188.930** | **-4.166** | **4.70E-05*** ** |
| Map Task: Fixation | **-0.065** | **0.021** | **181.751** | **-3.123** | **0.002** ** |
| ROI Task: Fixation * Map Task: Fixation | **0.155** | **0.030** | **181.751** | **5.117** | **7.85E-07*** ** |
| **Adults** | | | | | |
| Intercept (ROI Task: One-back \| Map Task: One-back) | **0.198** | **0.013** | **32.895** | **15.501** | **1.11E-16*** ** |
| ROI Task: Fixation | **-0.041** | **0.014** | **36.000** | **-2.918** | **6.03E-03*** ** |
| Map Task: Fixation | -0.019 | 0.014 | 36.000 | -1.332 | 0.191 |
| ROI Task: Fixation * Map Task: Fixation | **0.051** | **0.020** | **36.000** | **2.567** | **0.015*** |

**Table S5 | LME Results for Text Selectivity by Split Task VWFA on Split Task Data**
Linear mixed effects model results for text-selectivity index prediction as a function of the two way interaction between ROI task and BOLD task in children (**a**) and adult (**b**) participants.. Significant results are displayed in bold and asterisks indicate the degree of significance ($p < 0.001$: ***, $p < 0.01$: **, $p < 0.05$: *)

| Visual Word Form Area and Fusiform Face Area Size Differences | | | | | | |
|---|---|---|---|---|---|---|
| | **t** | **CI** | | **DOF** | ***p*** | |
| | *(VWFA > FFA)* | *low* | *high* | | | |
| Children | **-11.685** | **-831.621** | **-589.623** | **81** | **4.65E-19** | *** |
| Adults | **-2.614** | **-727.199** | **-69.087** | **13** | **0.021** | * |

**Table S6 | Visual Word Form Area and Fusiform Face Area Size Differences**
Results from a two-tailed, paired-samples t-test comparing the size of the Visual Word Form Area (VWFA; in $\log_{10}$ number of vertices) to the size of the Fusiform Face Area (FFA) in child and adult participants. Significant results are displayed in bold and asterisks indicate the degree of significance (p < 0.001: ***, p < 0.01: **, p < 0.05: *)

| | Individual Medoid to Group Medoid Distances | | | | | | | | |
|---|---|---|---|---|---|---|---|---|---|
| | VWFA | | FFA | | t | CI | | DOF | *p* |
| | *Mean* | *SD* | *Mean* | *SD* | *(VWFA > FFA)* | *low* | *high* | | |
| Children | **13.904** | **10.024** | **10.919** | **8.595** | **4.714** | **3.470** | **8.471** | **162** | **5.20E-06** *** |
| Adults | 10.125 | 7.896 | 9.040 | 6.754 | 0.830 | -3.205 | 7.545 | 26 | 0.414 |

**Table S7 | Individual Medoid to Group Medoid Distances**
Average distance from the individual region of interest (ROI) medoid (center of mass) to the group ROI medoid for both the Visual Word Form Area (VWFA) and the Fusiform Face Area (FFA) in mm. Results from a two-tailed, paired-samples t-test comparing the distances from VWFA to the distances from FFA are also displayed in child and adult participants. Significant results are displayed in bold and asterisks indicate the degree of significance ($p < 0.001$: ***, $p < 0.01$: **, $p < 0.05$: *)

**a**

| BOLD Response ~ ROI + (1 \| Participant) - Children | β | Std. Err | DOF | t | p |
|---|---|---|---|---|---|
| **Text** | | | | | |
| Intercept (native VWFA) | **1.048** | **0.055** | **228.980** | **19.191** | **1.43E-49***** |
| cVWFA | **-0.154** | **0.055** | **313.070** | **-2.821** | **0.005**** |
| aVWFA | **-0.470** | **0.055** | **313.070** | **-8.602** | **3.83E-16***** |
| rVWFA | **-0.187** | **0.055** | **313.070** | **-3.420** | **0.001***** |
| kVWFA | -0.011 | 0.055 | 313.070 | -0.201 | 0.841 |
| **Pseudofonts** | | | | | |
| Intercept (native VWFA) | **0.704** | **0.051** | **254.577** | **13.832** | **7.84E-33***** |
| cVWFA | 0.089 | 0.053 | 313.509 | 1.667 | 0.096 |
| aVWFA | **-0.178** | **0.053** | **313.509** | **-3.335** | **0.001***** |
| rVWFA | **0.161** | **0.053** | **313.509** | **3.017** | **0.003**** |
| kVWFA | **0.280** | **0.053** | **313.509** | **5.251** | **2.80E-07***** |
| **Faces** | | | | | |
| Intercept (native VWFA) | **0.247** | **0.061** | **171.889** | **4.067** | **7.23E-05***** |
| cVWFA | **0.526** | **0.052** | **311.589** | **10.043** | **9.59E-21***** |
| aVWFA | **0.267** | **0.052** | **311.589** | **5.089** | **6.22E-07***** |
| rVWFA | **0.566** | **0.052** | **311.589** | **10.811** | **2.41E-23***** |
| kVWFA | **1.108** | **0.052** | **311.589** | **21.146** | **3.56E-62***** |
| **Objects** | | | | | |
| Intercept (native VWFA) | **0.592** | **0.068** | **178.579** | **8.708** | **2.04E-15***** |
| cVWFA | **0.547** | **0.060** | **311.444** | **9.106** | **1.06E-17***** |
| aVWFA | **0.223** | **0.060** | **311.444** | **3.712** | **2.44E-04***** |
| rVWFA | **0.757** | **0.060** | **311.444** | **12.605** | **1.02E-29***** |
| kVWFA | **0.983** | **0.060** | **311.444** | **16.386** | **5.97E-44***** |
| **Limbs** | | | | | |
| Intercept (native VWFA) | **0.622** | **0.069** | **173.337** | **8.956** | **5.14E-16***** |
| cVWFA | **0.600** | **0.060** | **311.184** | **9.947** | **2.02E-20***** |
| aVWFA | **0.288** | **0.060** | **311.184** | **4.767** | **2.87E-06***** |
| rVWFA | **0.773** | **0.060** | **311.184** | **12.810** | **1.84E-30***** |
| kVWFA | **1.029** | **0.060** | **311.184** | **17.051** | **1.70E-46***** |

**b**

| BOLD Response ~ ROI + (1 \| Participant) - Adults | | | | | |
|---|---|---|---|---|---|
| | **β** | **Std. Err** | **DOF** | **t** | **p** |
| **Text** | | | | | |
| Intercept (native VWFA) | **1.182** | **0.128** | **28.981** | **9.237** | **3.89E-10*** ** |
| cVWFA | **-0.259** | **0.117** | **51.151** | **-2.217** | **0.031*** |
| aVWFA | **-0.514** | **0.117** | **51.151** | **-4.390** | **5.71E-05*** ** |
| rVWFA | **-0.250** | **0.117** | **51.151** | **-2.140** | **0.037*** |
| kVWFA | -0.194 | 0.117 | 51.151 | -1.653 | 0.104 |
| **Pseudofonts** | | | | | |
| Intercept (native VWFA) | **0.770** | **0.114** | **23.932** | **6.783** | **5.21E-07*** ** |
| cVWFA | 0.072 | 0.092 | 51.114 | 0.791 | 0.433 |
| aVWFA | **-0.188** | **0.092** | **51.114** | **-2.050** | **0.046*** |
| rVWFA | 0.139 | 0.092 | 51.114 | 1.523 | 0.134 |
| kVWFA | 0.157 | 0.092 | 51.114 | 1.715 | 0.092 |
| **Faces** | | | | | |
| Intercept (native VWFA) | **0.390** | **0.097** | **31.825** | **4.037** | **3.18E-04*** ** |
| cVWFA | **0.372** | **0.093** | **51.216** | **4.004** | **2.01E-04*** ** |
| aVWFA | 0.105 | 0.093 | 51.216 | 1.130 | 0.264 |
| rVWFA | **0.542** | **0.093** | **51.216** | **5.846** | **3.53E-07*** ** |
| kVWFA | **0.831** | **0.093** | **51.216** | **8.954** | **4.68E-12*** ** |
| **Objects** | | | | | |
| Intercept (native VWFA) | **0.488** | **0.108** | **24.963** | **4.519** | **1.30E-04*** ** |
| cVWFA | **0.443** | **0.090** | **51.139** | **4.939** | **8.79E-06*** ** |
| aVWFA | 0.162 | 0.090 | 51.139 | 1.807 | 0.077 |
| rVWFA | **0.698** | **0.090** | **51.139** | **7.779** | **3.17E-10*** ** |
| kVWFA | **0.739** | **0.090** | **51.139** | **8.241** | **5.99E-11*** ** |
| **Limbs** | | | | | |
| Intercept (native VWFA) | **0.552** | **0.102** | **28.405** | **5.394** | **9.04E-06*** ** |
| cVWFA | **0.530** | **0.092** | **51.188** | **5.743** | **5.11E-07*** ** |
| aVWFA | **0.227** | **0.092** | **51.188** | **2.462** | **0.017*** |
| rVWFA | **0.727** | **0.092** | **51.188** | **7.880** | **2.19E-10*** ** |
| kVWFA | **0.761** | **0.092** | **51.188** | **8.241** | **5.94E-11*** ** |

**Table S8 | Group and Template Visual Word Form Area Activation LME Results**
Linear mixed effects model results for BOLD response prediction as a function of the Visual Word Form Area (VWFA) for each stimulus category in child (**a**) and adult (**b**) participants. Manually-defined, native VWFAs are treated as the reference region compared with the child (cVWFA) and adult (aVWFA) participants' probabilistic VWFA, along with two literature-based template VWFAs (rVWFA from Rosenke et al., 2018 and kVWFA from Kubota et al., 2023). Significant results are displayed in bold and asterisks indicate the degree of significance ($p < 0.001$: ***, $p < 0.01$: **, $p < 0.05$: *).

**a**

| BOLD Response ~ ROI + (1 \| Participant) - Children | | | | | |
|---|---|---|---|---|---|
| | **β** | **Std. Err** | **DOF** | **t** | **p** |
| **Text** | | | | | |
| Intercept (native FFA) | **0.624** | **0.038** | **110.782** | **16.403** | **2.05E-31*** ** |
| cFFA | **0.166** | **0.023** | **324.000** | **7.187** | **4.60E-12*** ** |
| aFFA | **0.282** | **0.023** | **324.000** | **12.174** | **2.47E-28*** ** |
| rFFA | 0.011 | 0.023 | 324.000 | 0.465 | 0.642 |
| kFFA | 0.037 | 0.023 | 324.000 | 1.585 | 0.114 |
| **Pseudofonts** | | | | | |
| Intercept (native FFA) | **0.540** | **0.034** | **118.693** | **16.064** | **1.50E-31*** ** |
| cFFA | **0.188** | **0.022** | **324.000** | **8.376** | **1.68E-15*** ** |
| aFFA | **0.312** | **0.022** | **324.000** | **13.868** | **1.18E-34*** ** |
| rFFA | 0.013 | 0.022 | 324.000 | 0.597 | 0.551 |
| kFFA | 0.038 | 0.022 | 324.000 | 1.670 | 0.096 |
| **Faces** | | | | | |
| Intercept (native FFA) | **1.704** | **0.047** | **120.971** | **36.130** | **1.16E-66*** ** |
| cFFA | **-0.285** | **0.032** | **324.000** | **-8.848** | **5.90E-17*** ** |
| aFFA | **-0.192** | **0.032** | **324.000** | **-5.948** | **7.02E-09*** ** |
| rFFA | **-0.629** | **0.032** | **324.000** | **-19.489** | **1.58E-56*** ** |
| kFFA | **-0.416** | **0.032** | **324.000** | **-12.894** | **5.49E-31*** ** |
| **Objects** | | | | | |
| Intercept (native FFA) | **1.319** | **0.050** | **112.879** | **26.142** | **1.05E-49*** ** |
| cFFA | **0.216** | **0.032** | **324.000** | **6.844** | **3.87E-11*** ** |
| aFFA | **0.370** | **0.032** | **324.000** | **11.711** | **1.19E-26*** ** |
| rFFA | **-0.192** | **0.032** | **324.000** | **-6.087** | **3.25E-09*** ** |
| kFFA | 0.024 | 0.032 | 324.000 | 0.768 | 0.443 |
| **Limbs** | | | | | |
| Intercept (native FFA) | **1.404** | **0.049** | **108.686** | **28.372** | **4.61E-52*** ** |
| cFFA | **0.110** | **0.029** | **324.000** | **3.760** | **2.02E-04*** ** |
| aFFA | **0.271** | **0.029** | **324.000** | **9.268** | **2.74E-18*** ** |
| rFFA | **-0.206** | **0.029** | **324.000** | **-7.037** | **1.17E-11*** ** |
| kFFA | -0.044 | 0.029 | 324.000 | -1.490 | 0.137 |

**b**

| BOLD Response ~ ROI + (1 \| Participant) - Adults | β | Std. Err | DOF | t | p |
|---|---|---|---|---|---|
| **Text** | | | | | |
| Intercept (native FFA) | **0.797** | **0.083** | **26.992** | **9.602** | **3.38E-10*** ** |
| cFFA | 0.020 | 0.075 | 52.000 | 0.266 | 0.791 |
| aFFA | 0.124 | 0.075 | 52.000 | 1.653 | 0.104 |
| rFFA | -0.140 | 0.075 | 52.000 | -1.865 | 0.068 |
| kFFA | -0.109 | 0.075 | 52.000 | -1.454 | 0.152 |
| **Pseudofonts** | | | | | |
| Intercept (native FFA) | **0.725** | **0.081** | **24.152** | **8.905** | **4.26E-09*** ** |
| cFFA | 0.054 | 0.068 | 52.000 | 0.797 | 0.429 |
| aFFA | **0.162** | **0.068** | **52.000** | **2.375** | **0.021*** |
| rFFA | **-0.153** | **0.068** | **52.000** | **-2.247** | **0.029*** |
| kFFA | -0.095 | 0.068 | 52.000 | -1.392 | 0.170 |
| **Faces** | | | | | |
| Intercept (native FFA) | **1.770** | **0.095** | **33.104** | **18.607** | **4.23E-19*** ** |
| cFFA | **-0.567** | **0.096** | **52.000** | **-5.905** | **2.72E-07*** ** |
| aFFA | **-0.402** | **0.096** | **52.000** | **-4.185** | **1.10E-04*** ** |
| rFFA | **-0.825** | **0.096** | **52.000** | **-8.598** | **1.47E-11*** ** |
| kFFA | **-0.682** | **0.096** | **52.000** | **-7.108** | **3.32E-09*** ** |
| **Objects** | | | | | |
| Intercept (native FFA) | **1.321** | **0.123** | **22.420** | **10.718** | **2.72E-10*** ** |
| cFFA | -0.069 | 0.097 | 52.000 | -0.706 | 0.483 |
| aFFA | 0.079 | 0.097 | 52.000 | 0.817 | 0.418 |
| rFFA | **-0.422** | **0.097** | **52.000** | **-4.344** | **6.50E-05*** ** |
| kFFA | **-0.237** | **0.097** | **52.000** | **-2.439** | **0.018*** |
| **Limbs** | | | | | |
| Intercept (native FFA) | **1.359** | **0.106** | **23.630** | **12.850** | **3.76E-12*** ** |
| cFFA | -0.162 | 0.087 | 52.000 | -1.866 | 0.068 |
| aFFA | -0.001 | 0.087 | 52.000 | -0.016 | 0.987 |
| rFFA | **-0.438** | **0.087** | **52.000** | **-5.040** | **5.98E-06*** ** |
| kFFA | **-0.333** | **0.087** | **52.000** | **-3.830** | **3.47E-04*** ** |

**Table S9 | Group and Template Fusiform Face Area Activation LME Results**
Linear mixed effects model results for BOLD response prediction as a function of the Fusiform Face Area (FFA) for each stimulus category in child (**a**) and adult (**b**) participants. Manually-defined, native FFAs are treated as the reference region compared with the child (cFFA) and adult (aFFA) participants' probabilistic FFA, along with two literature-based template FFAs (rFFA from Rosenke et al., 2018 and kFFA from Kubota et al., 2023). Significant results are displayed in bold and asterisks indicate the degree of significance (p < 0.001: ***, p < 0.01: **, p < 0.05: *)

| Text Selectivity ~ ROI + (1 \| Participant) - Adults | | | | | | |
|---|---|---|---|---|---|---|
| | β | Std. Err | DOF | t | p | |
| **Children** | | | | | | |
| Intercept (native VWFA) | **0.160** | **0.009** | **219.427** | **18.733** | **2.09E-47** | *** |
| cVWFA | **-0.185** | **0.008** | **312.034** | **-22.036** | **1.48E-65** | *** |
| aVWFA | **-0.193** | **0.008** | **312.034** | **-22.971** | **4.77E-69** | *** |
| rVWFA | **-0.222** | **0.008** | **312.034** | **-26.437** | **1.21E-81** | *** |
| kVWFA | **-0.240** | **0.008** | **312.034** | **-28.600** | **3.40E-89** | *** |
| **Adults** | | | | | | |
| Intercept (native VWFA) | **0.174** | **0.018** | **39.637** | **9.849** | **3.29E-12** | *** |
| cVWFA | **-0.184** | **0.019** | **51.209** | **-9.798** | **2.46E-13** | *** |
| aVWFA | **-0.168** | **0.019** | **51.209** | **-8.926** | **5.16E-12** | *** |
| rVWFA | **-0.213** | **0.019** | **51.209** | **-11.302** | **1.59E-15** | *** |
| kVWFA | **-0.222** | **0.019** | **51.209** | **-11.779** | **3.41E-16** | *** |

**Table S10 | Group and Template Visual Word Form Area Text Selectivity LME Results**
Linear mixed effects model results for text-selectivity index prediction as a function of the Visual Word Form Area (VWFA) in child and adult participants. Manually-defined, native VWFAs are treated as the reference region compared with the child (cVWFA) and adult (aVWFA) participants' probabilistic VWFA, along with two literature-based template VWFAs (rVWFA from Rosenke et al., 2018 and kVWFA from Kubota et al., 2023). Significant results are displayed in bold and asterisks indicate the degree of significance ($p < 0.001$: ***, $p < 0.01$: **, $p < 0.05$: *)
